# seqLens: optimizing language models for genomic predictions

**DOI:** 10.1101/2025.03.12.642848

**Authors:** Mahdi Baghbanzadeh, Brendan Mann, Keith A. Crandall, Ali Rahnavard

**Author notes:** Contributing authors.

## Abstract

Understanding evolutionary variation in genomic sequences through the lens of language modeling has the potential to revolutionize biological research. Yet to maximize the utility of language modeling in genomics, we must overcome computational challenges in tokenization and model architecture adapted to diverse genomic features across evolutionary timescales. In this study, we investigated key elements in genomic language modeling (gLM), including tokenization, pretraining datasets, fine-tuning approaches, pooling methods, and domain adaptation, and applied the language models to diverse genomic data. We gathered two evolutionarily distinct pretraining datasets: one consisting of 19,551 reference genomes, including over 18,000 prokaryotic genomes (115B nucleotides) and the remainder eukaryotic genomes, and another more balanced dataset with 1,354 genomes, including 1,166 prokaryotic and 188 eukaryotic reference genomes (180B nucleotides). We trained five byte-pair encoding tokenizers and pretrained 52 gLMs, systematically comparing different architectures, hyperparameters, and classification heads. We introduce `seqLens`, a family of models based on disentangled attention with relative positional encoding, which outperforms relatively similar-sized models in 13 of 19 benchmarking phenotypic predictions. We further explore continual pretraining, domain adaptation, and parameter-efficient fine-tuning methods to assess trade-offs between computational efficiency and accuracy. Our findings demonstrate that relevant pretraining data significantly boost performance, alternative pooling techniques can enhance classification, tokenizers with larger vocabulary sizes negatively impact generalization, and gLMs are capable of understanding evolutionary relationships. These insights provide a foundation for optimizing genomic language models for identifying diverse evolutionary genomic features and improving genome annotations.

## 1 Main

DNA is a language [1–3] with many unexplored aspects due to the challenges associated with experimentally validating the function of genetic elements on a genome-wide scale and the diversity of function across evolutionary timescales [4]. Genomics and metagenomics have transformed data availability primarily due to rapid advances in high-throughput sequencing (HTS) technologies [5, 6]. These technologies have catalyzed the era of big data in genomics, generating vast amounts of information with precision and detail across the tree of life [7–9]. Improvements in HTS technologies have increased accessibility and cost-effectiveness in sequencing nucleic acids from across evolutionary diversity [10, 11]. However, a remaining challenge lies in extracting meaningful insights from this data deluge in a computationally efficient manner, which is where machine learning (ML) and deep learning (DL) techniques have emerged as invaluable tools [12–15].

The application of DL to DNA sequences has evolved from task-specific implementations to more generalized approaches using language models. Early studies explored neural network implementations for various tasks, including splice site prediction [16], modeling genetic responses to specific medications [17], identifying chromatin accessibility prediction [18], transcription factor binding sites [19], and performing taxonomic profiling [1, 20, 21]. In contrast to these studies, which focused on training separate models for specific tasks, DNA foundation models such as DNABERT [2], DNABERT-2 [3], GROVER [22], HyenaDNA [23], and Nucleotide Transformer [24] offer a more generalized approach by first pretraining on large-scale genomic sequences to develop a broad understanding of genomic patterns and structures. Once pretrained, a single model can be adapted to various specific tasks through fine-tuning, which eliminates the need to train separate models for each application. However, existing models face limitations in tokenization methods, architectural design, parameter efficiency, and domain adaptation capabilities that affect their performance across diverse genomic applications. Moreover, a critical question remains whether these models truly capture biological principles or merely statistical patterns in genomic sequences. While studies demonstrate that models can learn regulatory elements and sequence constraints [24], it is unclear how well they grasp complex evolutionary relationships like phylogenetic structures, given computational constraints [25].

In this study, we develop innovative genomic language models and prediction strategies using advanced encoder architectures and benchmark each step to provide a comprehensive roadmap for genomic language model development. To facilitate the understanding and decoding of DNA sequences, we employ DeBERTa-v2 [26] and ESM [27, 28] architectures and introduce `seqLens` models. We systematically investigate dynamic tokenization using byte-pair encoding (BPE), diverse pretraining datasets spanning multiple species, architectural features such as disentangled attention, parameter-efficient fine-tuning approaches, different pooling methods, and continual pretraining for domain adaptation (Figure 1). We also evaluate our models’ understanding of evolutionary relationships by analyzing the embeddings and comparing performance against existing genomic language models. Our comprehensive benchmarking demonstrates that these methodological advances significantly enhance genomic language model performance, with one of the `seqLens` models outperforming similar-sized models in 13 of 19 benchmarking tasks, establishing a robust framework for developing future genomic language models that are more effective, adaptable, and capable of addressing complex biological questions.

**Fig. 1:**
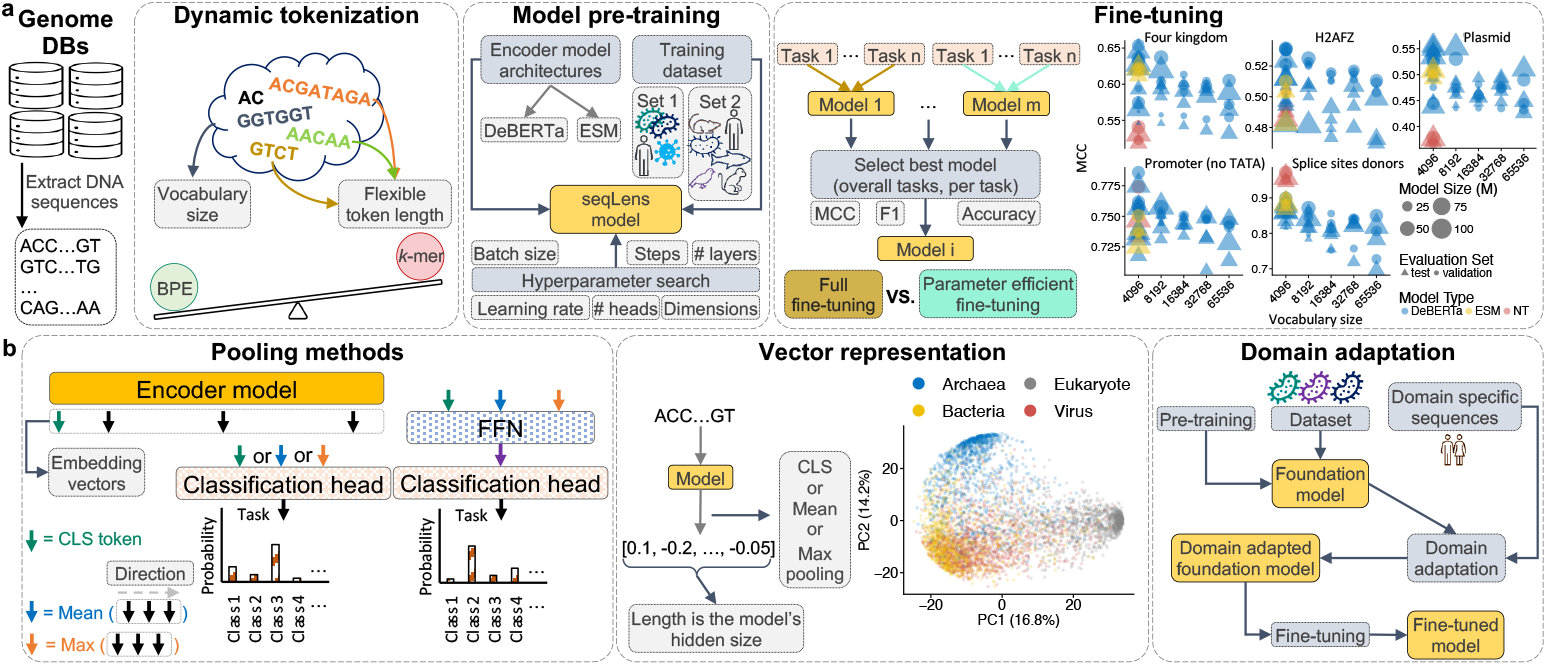
End-to-end DNA sequence modeling: core and auxiliary strategies. **a, first row**,The workflow begins with genome collection and dataset preparation, followed by BPE tokenization and model training using masked language modeling. Models are benchmarked on downstream tasks, and fine-tuning strategies are compared. **b, second row**, Various embedding pooling methods are evaluated for classification, along with genome vector representations and fine-tuning effects. Domain adaptation is also explored by continuing pretraining on a second dataset to assess performance impact.

## 2 Results

### 2.1 Byte-pair encoding tokenization improves convergence and performance in genomic language models

Tokenization is a crucial preliminary step in processing text sequences, as it defines how inputs are structured for models [29]. In our study, we selected the BPE approach to develop our tokenizers, a choice informed by established research in natural language processing [30] and previous investigations in applications of language modeling [31] in DNA [3, 22]. BPE is particularly suitable for handling large vocabularies and capturing subword information, making it a robust choice for diverse language data.

To comprehensively explore tokenizer behavior, we experimented with five different vocabulary sizes: 4,096, 8,192, 16,384, 32,768, and 65,536. We based our vocabulary size search on results reported in the DNABERT2 paper [3], where authors incrementally increased vocabulary size and found optimal performance at 4,096 tokens. We added more granularity to this analysis and investigated all benchmark datasets across the range from 2^12^ (4,096) to 2^16^ (65,536). We sampled 1 million sequences from the training dataset to train the tokenizers. We detail the key characteristics observed for each tokenizer after training in Table A1 and Figure A1a. For example, the 4096 tokenizer is like a dictionary with 4096 vocabularies, consisting of tokens such as {A, C, …, TCCTGGCC, TAAACCC}, with approximately 53% of them being *7* - mers (tokens of length 7) and the maximum length of tokens in this tokenizer is *10* -mers with frequency of 0.05% (Table A1). The varying maximum token lengths and frequency distributions highlight the trade-offs between vocabulary size and token representation efficiency. In general, larger vocabularies tend to produce longer tokens (Table A1), which can significantly impact the frequency distribution of token lengths, as illustrated in Figure A1. This behavior has a direct influence on the efficiency of model training and inference, affecting both computational costs and the model’s ability to understand language [32].

The BPE tokenizer’s ability to dynamically tokenize DNA sequences provides an edge in capturing complex data patterns, leading to better model performance, while the robust implementation ensures computational efficiency. The validation loss results (Figure A1b) demonstrate an advantage for the BPE tokenizer. When using the BPE tokenizer, both the `NT-50` and `NT-100` models achieved significantly lower validation losses compared to the scenario in which they utilized the fixed-length *6* -mer tokenizer. Furthermore, the models showed faster convergence with the BPE tokenizer, indicating a more efficient learning process. This suggests that the flexible, variable-length nature of the BPE tokenizer provides a better fit for the data, capturing more relevant patterns that aid the learning process. In terms of computational efficiency, the BPE tokenizer exhibited a higher TFLOP value compared to the fixed-length tokenizer, as illustrated in Figure A1c. It is also worth mentioning that we trained the models for 150K steps or a maximum time of 7 days. Since models with the *6* -mer tokenizer take longer to train, they did not reach the maximum number of steps in the given time. The results highlight the effectiveness of the BPE tokenizer in both model accuracy and training efficiency.

### 2.2 Smaller tokenizers lead to better generalization

We trained a set of models ranging from 15M to 111M parameters using various architectures, setups, and approaches (see Appendix B) to analyze a number of genomic tasks, including chromatin profiles, regulatory elements, genome origin, and splicing. There is no universally best-performing model (Figure C3), although `seqLens` models generally have a better score on test sets. Across datasets, models based on smaller vocabulary sizes generally outperform larger ones on the test set. As vocabulary size increases, model performance tends to decrease, although not uniformly (Figure 2a, Figure C4a, and Figure C5a). This can be attributed to two factors: the increased number of token embeddings requiring tuning and the reduced likelihood of tokens being sufficiently observed during pretraining. For example, for a given model with a *hidden dimension* of 512, each new vocabulary adds 512 new embedding weights to the model. So, the difference in parameter size of a model based on the 4,096 and the 65,536 tokenizers is more than (65536 − 4096) *>* 30*M* embedding weights. Also, if we consider a uniform chance of observing a token, the chances to train the weights of a token using the first tokenizer relative to the last tokenizer are roughly 65536*/*4096 *≈* 16 times more. Consequently, the weights of all tokens may not receive sufficient tuning due to the fixed number of training steps.

**Fig. 2:**
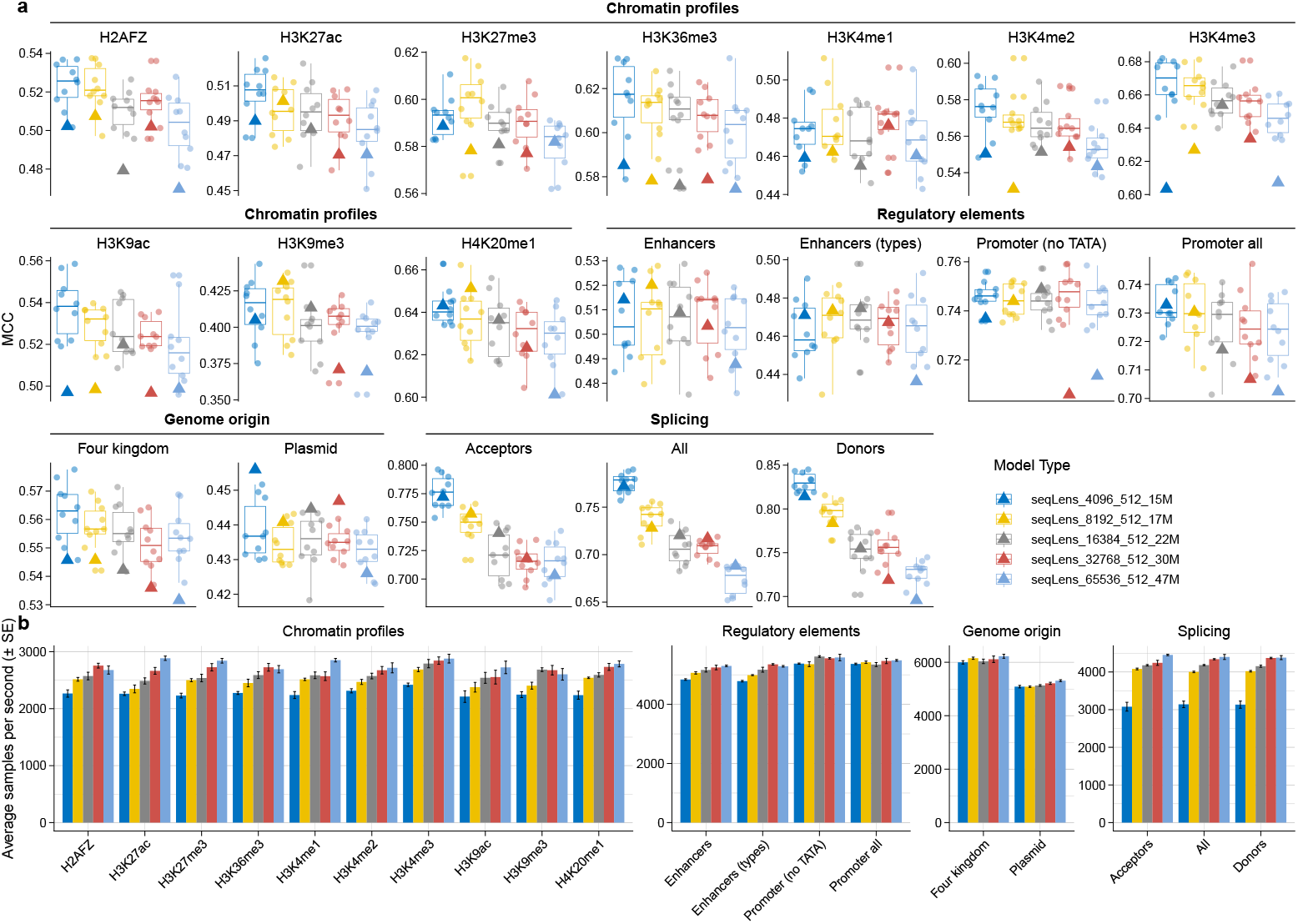
Validation and test scores of models across benchmarking datasets. **a**: The figure compares the performance of the models with the same parameter configuration but different tokenizer vocabulary sizes. All the models have 4 *layers*, 4 *attention heads, intermediate size* of 2048, and *hidden size* 512. Boxplots are based on the validation scores (circles), and the triangle represents the models’ performance on the test set. **b**: The figure compares the processing time of the models by comparing the average samples per second.

Larger tokenizers segment input sequences into longer tokens, resulting in shorter overall sequence lengths and, consequently, faster processing times (Figure 2b, Figure C4b, and Figure C5b). This effect is particularly noticeable in tasks such as *chromatin profiles*, where raw sequences are 1 Kbp in length, and *splicing* tasks, where the input sequences are 600 bp long. The reduction in sequence length due to larger tokenization units enhances computational efficiency, especially for longer genomic sequences.

### 2.3 Disentangled attention enhances seqLens model performance across most genomic tasks

One group of the `seqLens` models we trained involved varying parameter sizes of DeBERTa-v2 and an investigation into the effect of disentangled attention. Considering the performance on the test set, the `small` model outperforms its counterpart in 17 of 19 tasks, with exceptions in two *chromatin profiles* tasks (`H3K27me3` and `H3K4me1`) (Figure 3a). The `small` architecture differs from the other model with `46M` parameters by implementing disentangled attention and having 12 attention heads (instead of 6). Similarly, the `base` model outperforms the other model with `89M` parameters (its non-disentangled attention equivalent) in 15 of 19 tasks, with exceptions in three *chromatin profiles* tasks (`H3K4me1`, `H3K4me2`, and `H3K4me3`) and the `splicing acceptors` task (Figure 3). These results highlight the potential benefits of incorporating disentangled attention, particularly for most tasks, although certain *chromatin profiles* and *splicing* tasks remain challenging. Figure 3a also allows for a comparison among the `xsmall`, `small`, and `base` models. The `base` model demonstrates superior performance in 10 of 19 tasks, including the `four kingdoms` and `plasmid detection` datasets. The `xsmall` model performs better in 5 tasks, showcasing that smaller models can sometimes excel in specific scenarios.

**Fig. 3:**
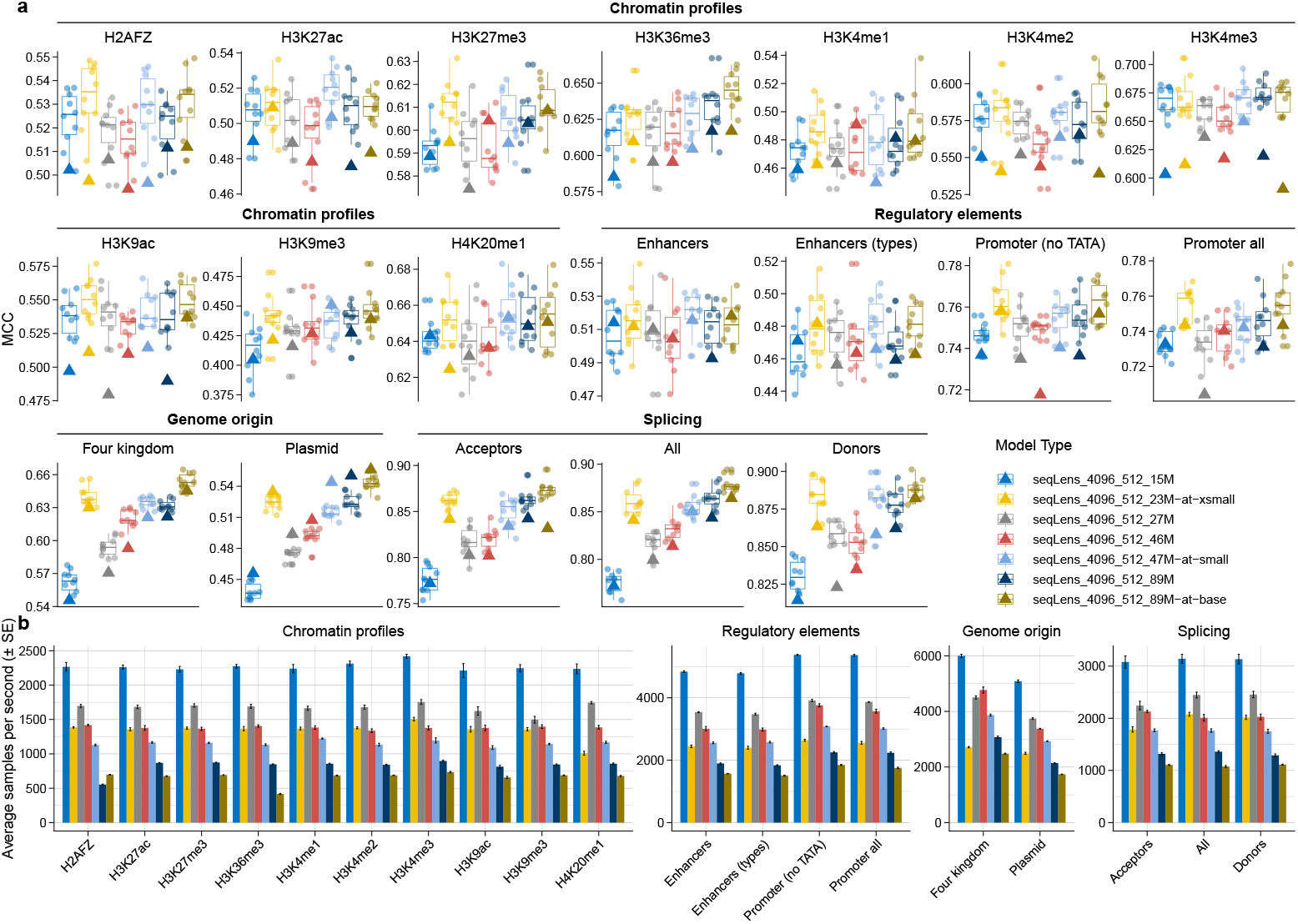
Benchmarking results of seqLens models with varying parameter sizes and the effect of disentangled attention. **a**: The figure compares the performance of seven `seqLens` models on 19 tasks, highlighting the impact of disentangled attention (models with `at` in their names). The boxes are based on the validation scores, and the test score is marked with a triangle. The `small` model outperforms its counterpart without disentangled attention in 17 of 19 tasks. Similarly, the `base` model outperforms its non-disentangled equivalent in 15 of 19 tasks. Comparisons among `xsmall`, `small`, and `base` models show that the `base` model performs best in 10 of 19 tasks. Boxplots are based on the validation scores (circles) and the triangle represents the models’ performance on the test set. **b**: The figure compares the processing time of the models by comparing the average samples per second.

With the same tokenizers, larger models process sequences more slowly than smaller models (Figure 3b). For example, the `15M` model processes samples, on average, 2.5 times faster than the `89M` model. Additionally, disentangled attention led to slower sample processing, with the `89M` model being, on average, about 20% faster than the `89M-at-base` model.

### 2.4 seqLens models achieve strong performance across genomic tasks

We trained two of the `seqLens` models with configurations similar to `NT-50` and `NT-100` but with a context length of 512. By maintaining similar training datasets, tokenizers, and infrastructures, this approach enables a more direct comparison of architectures. In both the `four kingdoms` and `plasmid detection` datasets, the ESM-based `seqLens` models outperformed the NT models, likely due to the influence of training data Figure 4. However, overall, `NT-100` outperformed its equivalent model in 15 tasks, and `NT-50` outperformed its equivalent in 13 tasks. Since the backbone of these models is the same, except for the tokenizer, context length, pretraining dataset, and training steps, these results underscore the importance of pretraining dataset composition and its interaction with model architecture and tokenizer design in determining downstream task performance.

**Fig. 4:**
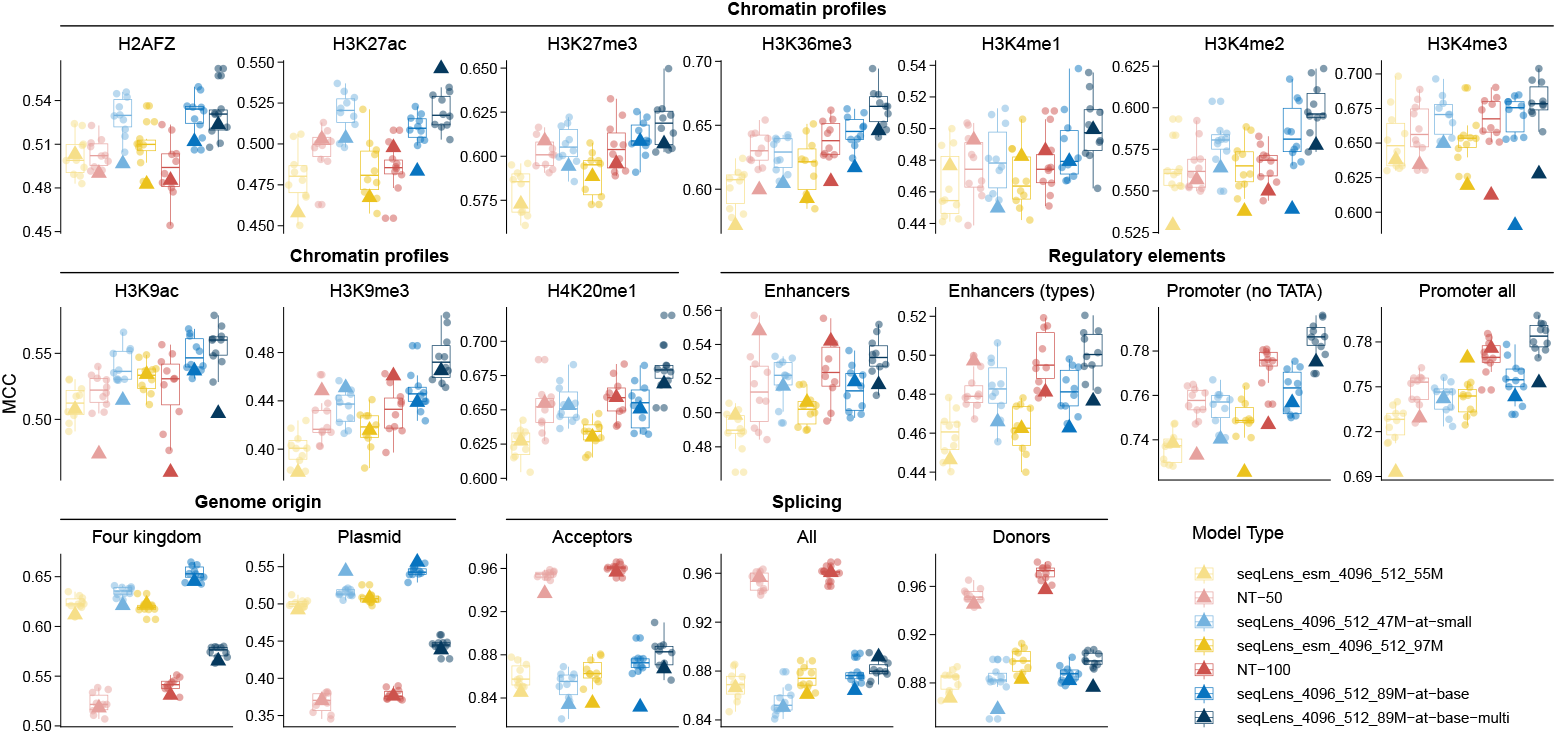
Comparison of validation and test scores for DeBERTa-inspired, ESM-based models, and NT models. The figure illustrates the performance of the DeBERTa-inspired models, NT models, and ESM-based models, which have similar configurations to `NT-50` and `NT-100` but are trained with the same setup as the DeBERTa-inspired models. Validation scores are represented as individual points within each box, while test scores are marked with triangles. Across 19 tasks, DeBERTa-inspired models (`small` and `base`) generally outperform ESM-based models. `base-multi` outperforms the other models in most cases. The results demonstrate that the DeBERTa-inspired models outperform ESM-based models, highlighting the importance of architectural choices and task-specific dynamics. Boxplots are based on the validation scores (circles), and the triangle represents the models’ performance on the test set.

Conversely, `47M-at-small` outperformed `seqLens esm 4096 512 55M` (similar to `NT-50`) in 14 of 19 tasks (Figure 4). Exceptions included two `chromatin profiles` tasks and all three `splicing` tasks. When comparing `47M-at-small` with `seqLens_esm_4096_512_97M` (similar to `NT-100`), which has nearly twice the parameters, the smaller model still outperformed the larger model in 12 of 19 tasks. Exceptions included two `chromatin profiles` tasks, all three `splicing` datasets, the `promoter all` dataset, and the `four kingdoms` dataset (where the performance difference was marginal: 0.621 vs. 0.622). For the larger models, `89M-at-base` outperformed `seqLens_esm_4096_512_55M` in 16 of 19 tasks. Exceptions were observed in the `H3K4me3` task and two `splicing` datasets (`acceptors` and `all`). Moreover, comparing `89M-at-base` with `seqLens_esm_4096_512_97M`, the former achieved better results in 12 of 19 tasks. Exceptions included two `chromatin profiles` tasks (`H3K4me1` and `H3K4me3`), the `promoter all` task, and two `splicing` datasets (`acceptors`: 0.832 vs. 0.835, and `donors`: 0.882 vs. 0.883). The only difference between these two types of models is their architecture. These results demonstrate the superiority of the DeBERTa-inspired `seqLens` models over the ESM models in generalizability and the ability to obtain better results in test datasets across benchmarking tasks.

Also, we trained a model similar to the configuration of the `89M-at-base` with an extended version of the NT pretraining datasets. This comparison will also help us understand the effect of the pretraining dataset. In 13 tasks `89M-at-base-multi` outperforms `NT-100` and `89M-at-base` and outperforms `NT-50` in 12 tasks (Figure 4). With this comparison, we can see the effect of pretraining dataset on the performance of the model on the downstream tasks.

These results highlight that the DeBERTa-inspired `seqLens` models often outperform ESM-based models across a majority of tasks despite having fewer parameters. The observed performance differences are not uniform, as ESM-based models excel in a few specific datasets, particularly those focused on splicing and chromatin profiles. This suggests that architectural differences and training dynamics, such as context length and parameter tuning, play critical roles in task-specific performance. Also, we observed that generally, the `seqLens` models with DeBERTa-v2 architecture provide better test scores across different tasks.

### 2.5 Max pooling achieves optimal unsupervised clustering performance for genomic sequence representations

UMAP [33] visualization of vector representations revealed distinct clustering patterns across different pooling methods and model states (Figure 5a,b). Fine-tuned models demonstrated significantly improved clustering and separation compared to pretrained models, with sequences of similar labels forming more cohesive groups in the reduced-dimensional space. The fine-tuned representations showed clear task-specific organization, while pretrained model vectors maintained more general, distributed patterns influenced by broader sequence characteristics.

**Fig. 5:**
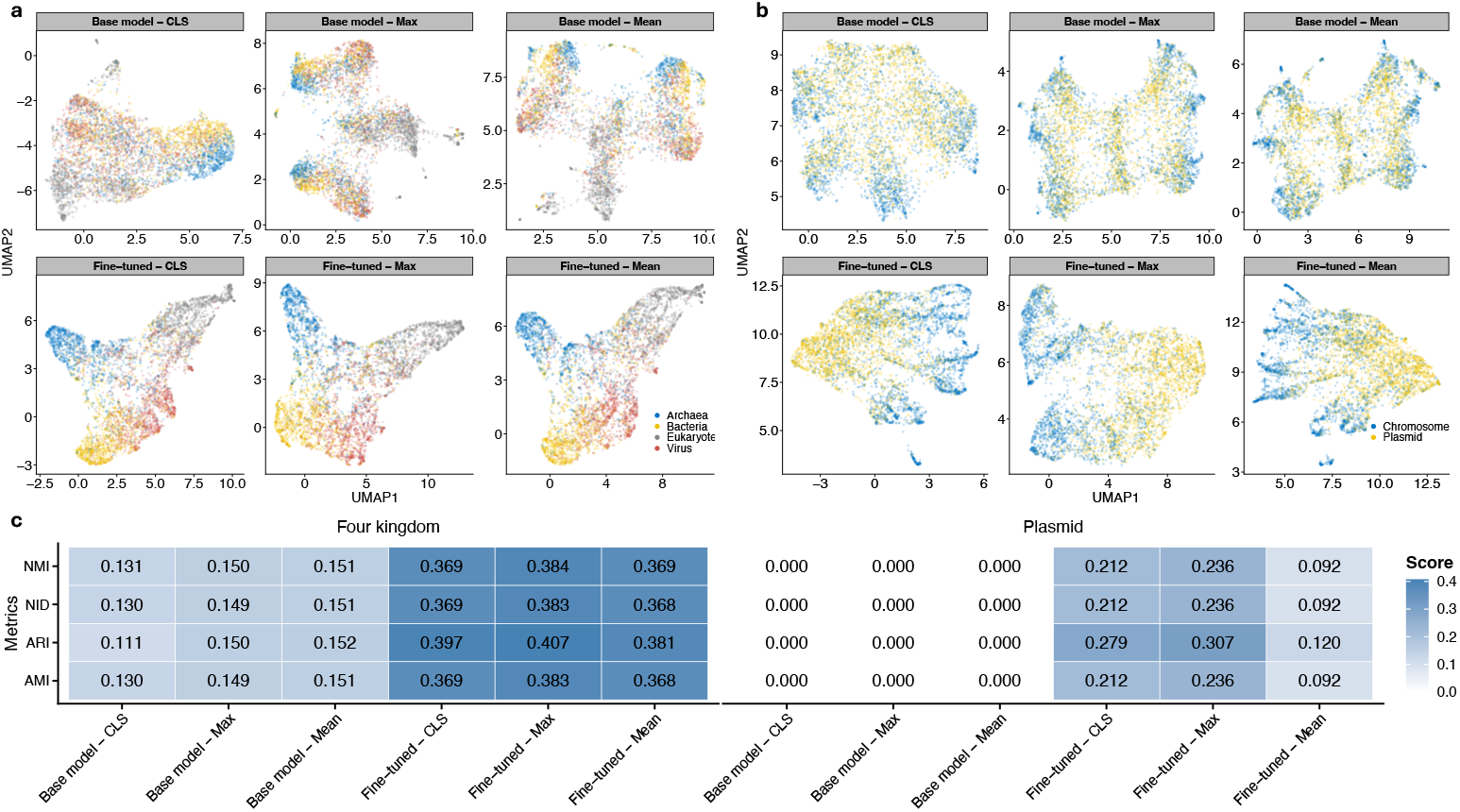
Analysis of vector representations from the base model on the four kingdoms and plasmid detection datasets. UMAP projections of vector representations for pretrained and fine-tuned models, summarized using three methods: `[CLS]` token, mean pooling, and max pooling. Fine-tuning leads to clearer clustering and better separation in both datasets. **a**, results for the `four kingdoms` dataset; **b**, results for the `plasmid detection` dataset; `four kingdoms` Clustering metric scores, adjusted rand index (ARI), adjusted mutual information (AMI), normalized information distance (NID), and normalized mutual information (NMI) for each summarization method on pretrained and fine-tuned models. Max pooling achieves the highest scores uniformly, indicating its superior ability to capture meaningful sequence representations.

Quantitative clustering analysis using adjusted Rand index (ARI), adjusted mutual information (AMI), normalized information distance (NID), and normalized mutual information (NMI) confirmed the visual observations (Figure 5c). Max pooling consistently achieved the highest scores across all clustering metrics in both datasets, demonstrating superior alignment between unsupervised clustering and true sequence labels. This performance advantage was maintained across both pretrained and fine-tuned model states, indicating that max pooling captures more discriminative sequence features regardless of model training state.

Dataset-specific patterns emerged in the clustering analysis. The `four kingdoms` dataset showed detectable clustering signal even with pretrained model representations, with vectors naturally separating according to taxonomic boundaries. In contrast, the `plasmid detection` dataset exhibited no meaningful clustering structure in pretrained representations (Figure 5c). This difference suggests that pretrained models inherently encode taxonomic relationships at higher phylogenetic levels but do not capture the subtle genomic features that distinguish plasmids from chromosomal sequences. Notably, for the `plasmid detection` dataset, mean pooling outperformed other methods in the pretrained setting, highlighting the context-dependent effectiveness of different pooling strategies.

### 2.6 seqLens embeddings recover phylogenetic structure and evolutionary signals in 16S rRNA sequences in a zero-shot setting

K-means clustering of `89M-at-base-multi` embeddings broadly recapitulated genus-level phylogenetic relationships (Figure 6a). t-SNE visualization revealed that sequences from the same genus formed cohesive clusters (Figure 6a,b), with some genera displaying partial overlap consistent with known taxonomic challenges within *Enterobacteriaceae*. K-means clustering on t-SNE coordinates yielded moderate but significant agreement with genus labels for `89M-at-base-multi` (NMI = 0.695, ARS = 0.480), indicating that embeddings capture genus-level evolutionary structure. In comparison, the Nucleotide Transformer models showed lower performance: `NT-100` achieved NMI = 0.519 and ARS = 0.366, while `NT-50` achieved NMI = 0.469 and ARS = 0.261 (Figure 6a).

**Fig. 6:**
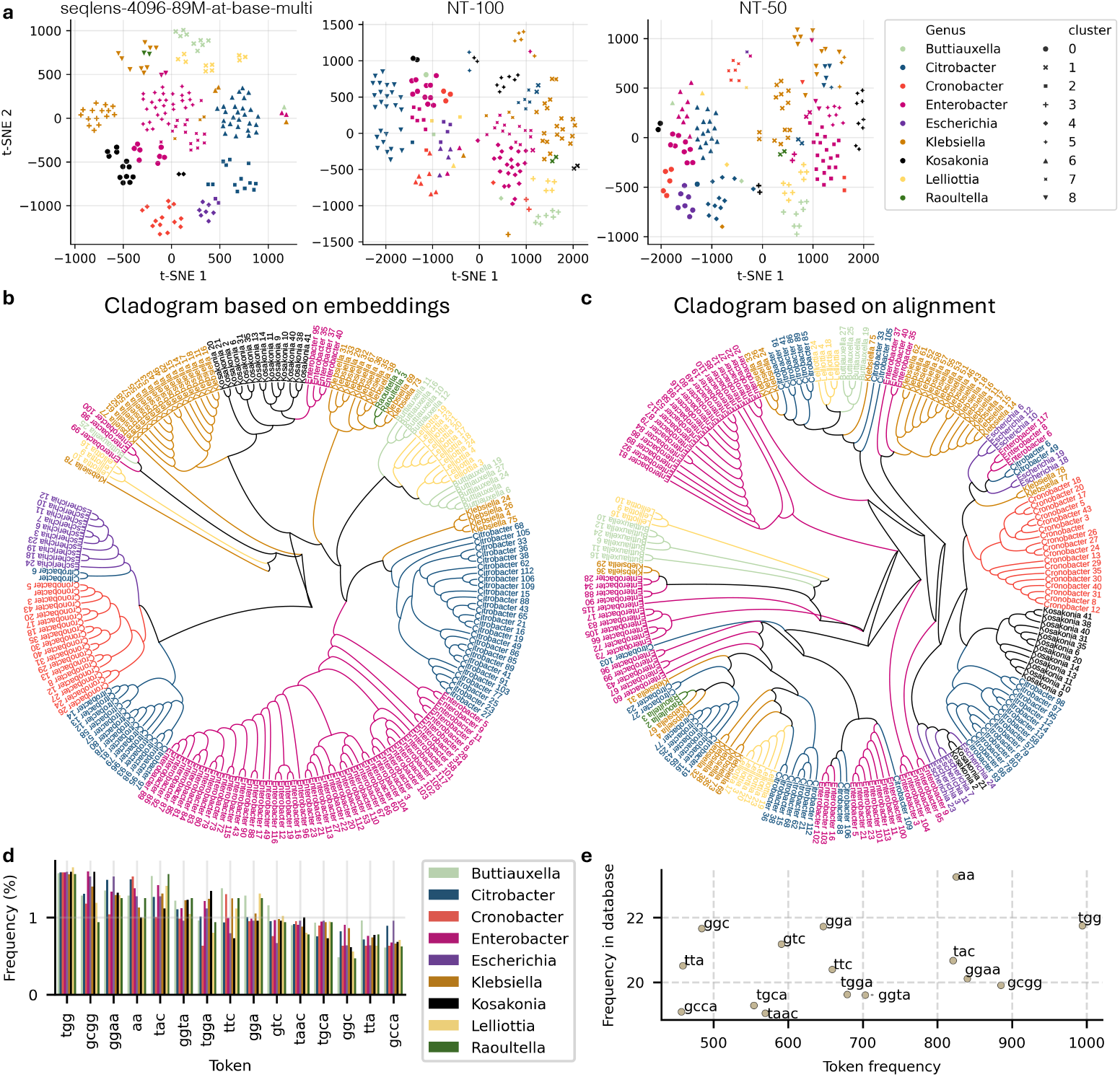
Comparison of seqlens embeddings with reference phylogeny. (a) Two-dimensional t-SNE projection of `seqlens`, and `NT` model embeddings, colored by genus and shaped by cluster assignment. (b) Cladogram reconstructed from embedding-derived distances (UPGMA). (c) Reference phylogeny reconstructed by maximum likelihood after multiple sequence alignment. (d) Relative frequencies of the top 15 tokens across genera, highlighting genus-specific differences. (e) Scatter plot of token frequency in `seqlens` (x-axis) versus motif frequency in the full 16S database (y-axis, log_2_ scale), showing divergence between learned tokens and raw sequence motif abundance.

Mantel correlation [34] between pairwise embedding distances (cosine) and patristic distances from the maximum likelihood phylogeny was statistically significant for `89M-at-base-multi` (Spearman *r* = 0.585, *p* = 0.001), demonstrating that embedding space preserves evolutionary distance relationships. The Nucleotide Transformer models showed weaker correlations: `NT-50` (Spearman *r* = 0.210, *p* = 0.001) and `NT-100` (Spearman *r* = 0.168, *p* = 0.001). A null model using randomly permuted embeddings showed no significant correlation (*r* = 0.007, *p* = 0.858), confirming that the observed correlation reflects genuine evolutionary signal rather than random structure.

Robinson-Foulds distance [35] between the embedding dendrogram (using `89M-at-base-multi`) and ML phylogeny indicated moderate topological agreement for `89M-at-base-multi` (RF = 300, normalized RF = 0.758), outperforming both `NT-50` (RF = 352, normalized RF = 0.889) and `NT-100` (RF = 342, normalized RF = 0.864). Monophyly analysis revealed genus-specific patterns: for `89M-at-base-multi`, *Raoultella* and *Cronobacter* were monophyletic in both trees (Figure 6b,c), while *Escherichia* achieved monophyly only in the embedding cladogram. In comparison, `NT-50` maintained monophyly only in *Raoultella*, while `NT-100` preserved monophyly in both *Cronobacter* and *Raoultella*. These results suggest that `89M-at-base-multi` may capture different and potentially more nuanced aspects of evolutionary relationships than both traditional sequence alignment-based methods and existing genomic language models.

Analysis of the 15 most frequent tokens revealed enrichment of motifs such as `TGG`, `GCGG`, and `GGAA` across genera, with notable genus-specific variation (Figure 6d,e). For instance, `GCGG` occurred at half the relative frequency in *Raoultella* compared to other genera, while `TGGA` was twice as frequent in *Kosakonia* as in *Cronobacter*. Importantly, token frequency after BPE tokenization did not correlate significantly with simple database frequency or fixed-length *k* -mer abundance (Pearson *r* = 0.299, *p* = 0.279; Spearman *ρ* = 0.336, *p* = 0.221; Kendall *τ* = 0.219, *p* = 0.281), indicating that the tokenizer identifies context-dependent sequence features rather than merely reflecting raw sequence frequency.

To test whether frequent tokens merely reflect conserved regions, we compared tokenizer output with motifs derived from the consensus sequence of all 200 aligned 16S genes. Remarkably, except for the top *3* -mer (`TTG`), none of the most common consensus-derived *3* -mers or *4* -mers appeared among the most frequent `seqLens` tokens.

### 2.7 Full parameter training outperforms LoRA in most tasks

LoRA implementation achieved substantial parameter reduction across both tested models, reducing trainable parameters to 0.65% (624,130 out of 96,406,373 parameters) for `NT-100` and 0.33% (296,450 out of 89,485,060 parameters) for `89M-at-base` using rank 8 and scaling factor 32. Notably, the results indicate that full parameter training consistently (17 of 19 tasks) outperforms LoRA fine-tuning across most scenarios (Figure 7). These performance differences may be attributed to several factors, including our specific choice of rank and scaling factor. Prior research has already suggested that LoRA fine-tuning might inherently lead to some performance reduction [36]. While LoRA is typically applied to large models (*>*1B parameters), here, we found that using it on smaller models (*∼*100M) did not improve performance.

**Fig. 7:**
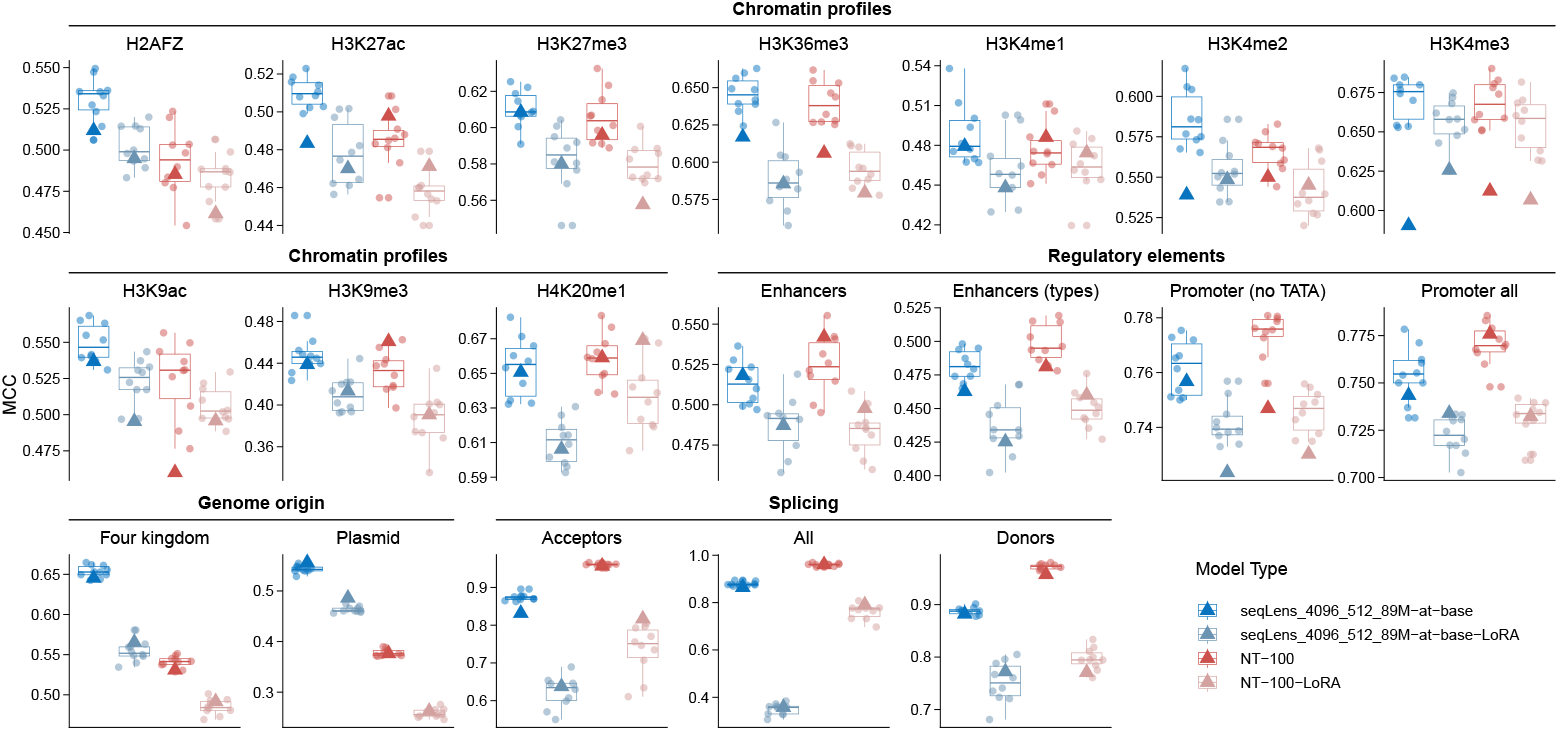
Performance comparison of full parameter training and LoRA fine-tuning. Each panel shows the results of the `NT-50` and `seqLens_4096_512_89M-at-base` models with two different approaches of full model and parameter efficient (LoRA) fine-tuning. Boxplots are based on the validation scores (circles), and the triangle represents the models’ performance on the test set.

### 2.8 Pooling strategy optimization reveals task-specific performance advantages

Classification performance varied significantly across pooling strategies, with the standard `[CLS]` token approach achieving best performance in 8 tasks, followed by `Mean` pooling (best in 6 tasks) and `Concat` (best in 5 tasks), while `Max` pooling demonstrated inferior performance overall (Figure 8a). `Concat` approach showed particular advantages in specific genomic tasks, excelling in `H3K36me3`, `promoter all`, `promoter (no TATA)`, `splice sites acceptors`, and `splice sites donors`. Despite the performance variations across methods, computational overhead analysis revealed no substantial differences in sample processing time between approaches (Figure 8b), indicating that the choice of pooling strategy can be optimized for task-specific performance without computational penalties. These results demonstrate that while `[CLS]` token remains effective for many classification tasks, alternative pooling strategies provide meaningful performance gains for specific genomic applications, warranting task-specific optimization of summarization methods.

**Fig. 8:**
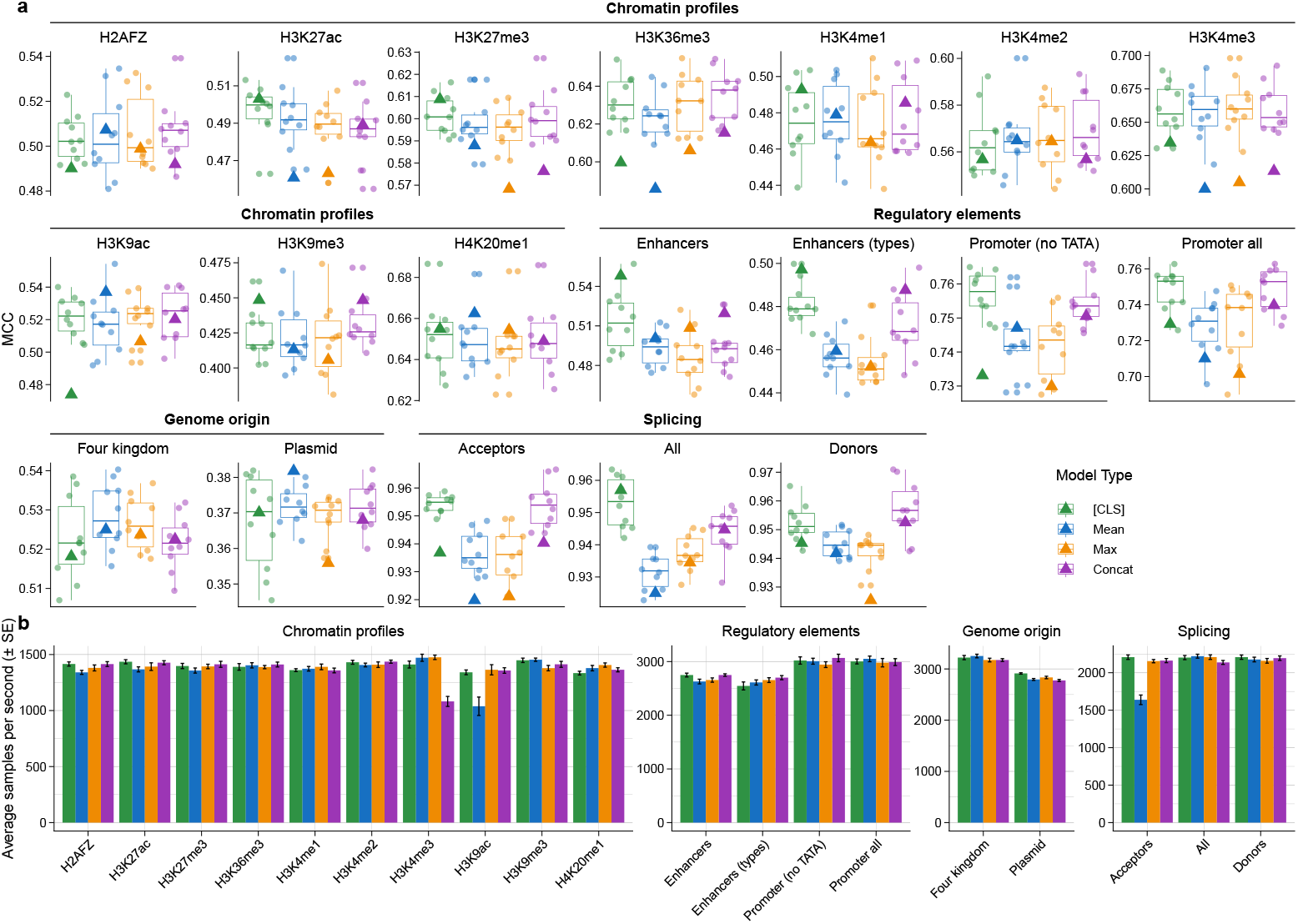
Comparison of classification performance across different fine-tuning strategies. **a**: The figure presents results for models using the `[CLS]` token, mean pooling (`Mean`), max pooling (`Max`), and the concatenated `CLS|Mean|Max` approach (`concat`). Boxplots are based on the validation scores (circles), and the triangle represents the models’ performance on the test set. **b**: The figure compares the processing time of the models by comparing the average samples per second.

### 2.9 Domain-adaptation boosts task-specific model performance

Domain-adaptive pretraining (DAP) demonstrated significant performance improvements with clear domain-specificity patterns. The eukaryote-adapted model (`Me`) achieved superior performance compared to the baseline `46M` model in 13 tasks and outperformed the general multi-species continued training (`Ms`) in 12 tasks (Figure 9a). However, `Me` showed reduced performance on broader taxonomic tasks, specifically showing lower performance in `four kingdoms` and `plasmid detection` compared to all other model variants. The prokaryote-adapted model (`Mp`) revealed the trade-offs inherent in domain specialization, with `Me` consistently outperforming `Mp` on human-centric tasks while showing inferior performance on prokaryote-related applications. Overall performance analysis showed `Me` achieving the best test results in 11 of 19 benchmarking tasks, establishing it as the top-performing model across the evaluation suite (Figure 9a). The prokaryote-adapted model (`Mp`) showed expected limitations, underperforming relative to the `46M` baseline and `Ms` on human-only genome and four kingdoms tasks (Figure 9a). However, `Mp` demonstrated domain-specific advantages in the `plasmid detection` task, outperforming both the `46M` baseline in validation and test sets and surpassing `Ms` in the test set. This performance pattern confirms the utility of prokaryote-specific adaptation for tasks closely aligned with the training domain (Figure 9a). Continued pretraining on the general multi-species dataset (`Ms`) improved performance over the `46M` baseline in 12 tasks, demonstrating that additional training on diverse genomic data enhances model capabilities without sacrificing versatility across taxonomically diverse applications (Figure 9a).

**Fig. 9:**
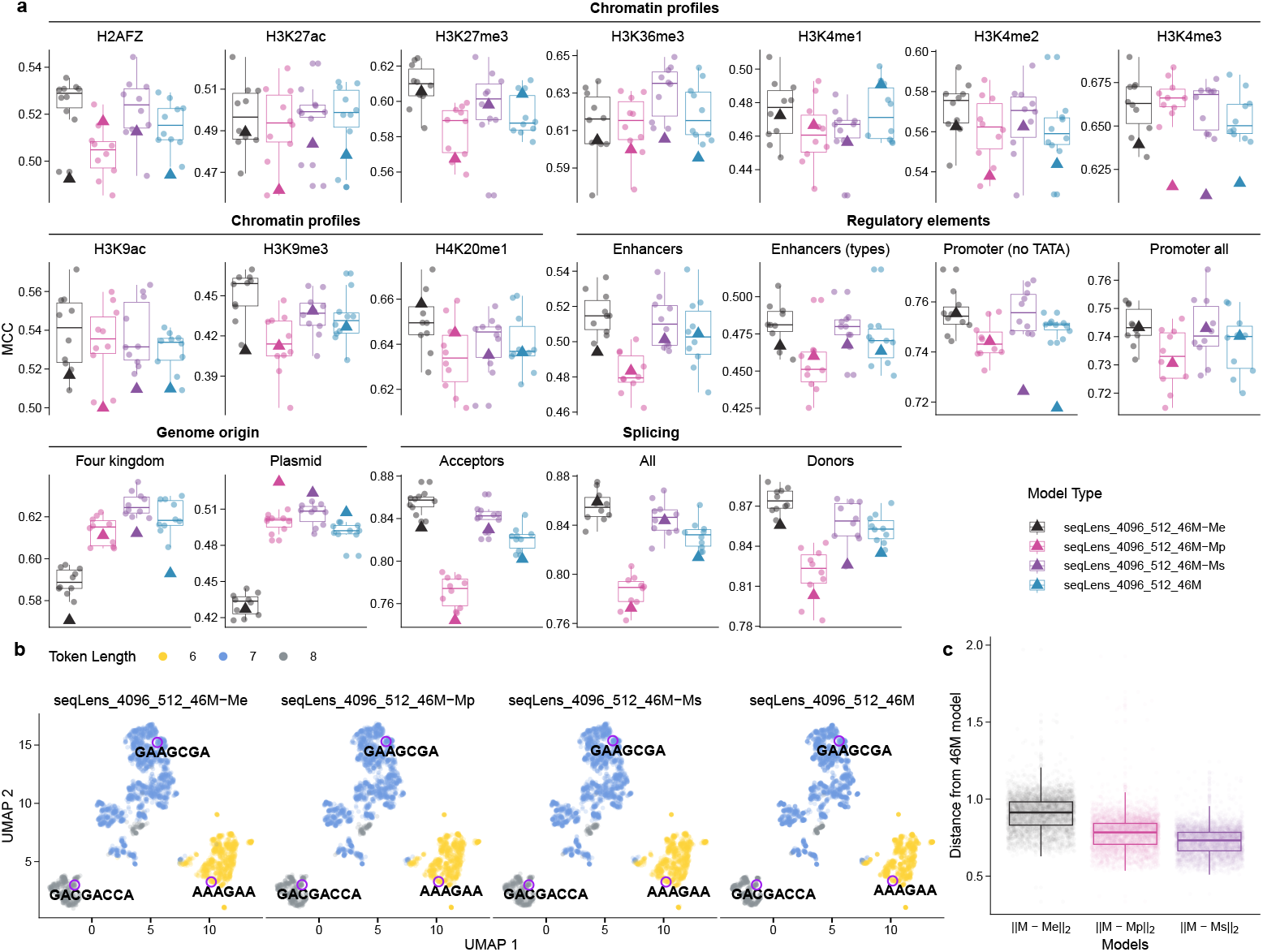
Effect of Domain-Adaptive Pretraining (DAP-training) on Model Performance Across Tasks. **a**: The `base-model` underwent an additional 150K training steps with three datasets: prokaryote-only genome (`Mp`), eukaryote-only genome (`Me`), and the original general-purpose training dataset (`Ms`). The resulting models were evaluated on benchmarking datasets. Boxplots are based on the validation scores (circles), and the triangle represents the models’ performance on the test set. **b**: The figure compares the 2d representation of token embeddings of models. Three sample tokens were marked to highlight the changes in representations. **c**: The distribution of token embedding distances from the `base-model`.

We also examined the token embeddings in the models and observed that the UMAP representations of token embeddings align with token lengths (Figure 9b). This suggests that token length plays a role in how embeddings are structured within the models and that models were able to detect it through training. Additionally, an analysis of the Euclidean distances between token embeddings of models from the `46M` model revealed that `Me` has the largest distance from `46M`, while `Ms` exhibits the most similar token embeddings to `46M` (Figure 9c). This suggests that exposing the model to a completely different domain dataset causes the largest shift in token embeddings compared to other scenarios.

## 3 Discussion

DNA sequences can be conceptualized as a complex language, rich with unseen patterns that hold the key to understanding fundamental biological insights [1–3]. Recent advancements in language modeling technology have revolutionized the ability to decipher these hidden patterns, offering faster, more efficient, and versatile methods compared to traditional alignment-based approaches [2, 3, 24]. Unlike conventional techniques that focus primarily on individual regions of DNA, language models provide a novel framework for exploring intricate interactions among regions, uncovering a deeper layer of genomic complexity. This paradigm shift aligns with the sequential nature of many biological techniques, such as mass spectrometry for protein analysis, suggesting that language models could play a transformative role in advancing our understanding of genomics [27, 28].

In this study, we investigated the impact of key features of language models on their performance in downstream genomic tasks. Our findings highlight several significant aspects that can enhance the design and utility of genomic language models. We demonstrated that a dynamic tokenizer, such as BPE, outperforms fixed-length *k* -mer-based tokenizers. Dynamic tokenization provides flexibility in capturing meaningful patterns and enables models to converge faster during training while achieving lower loss values [3]. This adaptability makes it a preferable choice for genomic language models. We also demonstrated that the BPE tokenizer processes text significantly faster than the native Python *k* -mer tokenizer. This speed-up can be attributed to the enhanced implementation efficiency of the BPE tokenizer, which leverages a high-performance library developed in Rust [37]. The Rust-based implementation is optimized for speed, enabling rapid tokenization and contributing to improved model efficiency. Despite the higher TFLOP requirements, this accelerated tokenization process likely plays a role in the superior performance observed with BPE-based models.

We also explored the DeBERTa-v2 architecture, particularly its implementation of disentangled attention, and compared it to equivalent models with the same parameter sizes. Our results show that architectures with disentangled attention consistently outperform their counterparts in most tasks and indicate a strong potential for scalability and effectiveness in larger implementations. We also investigated models with the same architecture but different vocabulary sizes. Our analysis revealed that models with smaller tokenizers tend to generalize better in downstream tasks. While a larger vocabulary can capture more nuanced linguistic patterns by introducing more distinct tokens, it also demands greater computational resources. Additionally, with a larger vocabulary, each token is seen less frequently on average during training, which may limit the model’s ability to learn robust representations [32].

Moreover, our findings highlight the importance of fine-tuning in refining vector representations for genomic data. While pretrained models provide general-purpose embeddings, fine-tuning enables the model to better capture task-specific distinctions, leading to improved clustering and separation of sequences. The differences observed between the `four kingdoms` and `plasmid detection` datasets suggest that pretrained embeddings inherently reflect broad genomic relationships but may lack the specificity needed for certain classification tasks. Additionally, the effectiveness of different pooling strategies underscores the need to carefully select summarization methods based on the nature of the task [38, 39]. We observed a contradiction between max pooling’s superior clustering performance and inferior classification accuracy. This can be explained by the fundamental difference between these evaluation paradigms. Max pooling creates more extreme, sparse representations by selecting only maximum activation values, which amplifies distinctive features [40] and creates sharper class boundaries in the embedding space—leading to better clustering metrics (ARI, AMI, NID, NMI). However, this aggressive feature selection discards potentially important contextual information and may result in representations that, while geometrically well-separated, are less robust for actual prediction tasks [41, 42].

The ability of genomic language models to capture evolutionary relationships without explicit phylogenetic training represents a significant advancement in computational biology [25]. Our results demonstrate that `seqLens` embeddings encode meaningful evolutionary relationships in a zero-shot manner, without requiring sequence alignment or explicit phylogenetic training, preserving evolutionary signal more effectively than existing Nucleotide Transformer models with superior correlation to phylogenetic distances and better topological agreement with maximum likelihood trees. The model’s ability to recover distance relationships while identifying genus-specific sequence patterns suggests potential applications in rapid taxonomic classification and evolutionary analysis of large-scale genomic datasets [25]. The divergence between language model-identified motifs and consensus-derived sequences demonstrates that the tokenizer captures recurrent but non-consensus motifs that may represent functionally important sequence variants or evolutionary intermediates, rather than simply recovering the most globally conserved alignment positions. However, the moderate topological agreement between embedding-based and traditional phylogenies raises questions about whether language model representations reflect true evolutionary history or alternative organizational principles based on functional or structural similarities. These findings suggest that genomic language models may complement traditional phylogenetic methods by providing novel perspectives on evolutionary relationships that capture both conserved evolutionary signal and functionally relevant sequence diversity, although further validation across diverse taxonomic groups and gene families is essential to establish their broader applicability in evolutionary genomics [43].

Additionally, we evaluated methods for leveraging language models in classification tasks. The conventional approach of using the `[CLS]` token from the last hidden state [44] does not consistently produce the best results. In fact, in more than half the cases, alternative methods such as mean pooling or concatenating the `[CLS]` token with mean and max pooling (`CLS|Mean|Max`) yield superior performance. This highlights the importance of exploring diverse strategies for summarizing model representations [45]. Using the [CLS] token or `Concat` approaches preserve more holistic sequence information and create balanced representations that generalize better to unseen data, explaining their superior classification performance despite lower clustering scores. This finding highlights that optimal clustering in embedding space does not necessarily translate to optimal predictive performance, particularly for complex DNA sequence analysis, where distributed patterns across multiple positions may be crucial [24]. We also investigated LoRA as a parameter-efficient fine-tuning method and found that while it reduces the computational costs of fine-tuning, it comes at the expense of model performance on downstream tasks compared to full parameter training. While the LoRA approach is highly beneficial for large language models [46] in terms of computing efficiency, our results indicate that for relatively small language models, the computational gains are not as significant as full model training is still achievable on a single GPU. Instead, we observe a loss of information that impacts overall model effectiveness. Future work could explore optimizing LoRA’s hyperparameters to minimize these performance gaps and better balance efficiency with accuracy.

The challenge of limited sample sizes in annotated samples [47] necessitates innovative approaches such as domain-adaptive pretraining (DAP), which adapts models to specific domains before fine-tuning [48, 49]. Our experiments with 150K training steps across prokaryote-only, eukaryote-only, and multi-species datasets reveal fundamental trade-offs between specialization and generalization: models adapted to narrow taxonomic domains excel within their target applications but suffer performance degradation on diverse tasks, while continued pretraining on diverse datasets maintains broader applicability with moderate improvements across various applications. These results demonstrate that DAP-training effectiveness depends critically on intended application scope, with narrow specialization suitable for focused research domains and diverse training preferable for general-purpose capabilities, thereby illustrating the significant potential of leveraging generic domain data to enhance model performance in niche applications where annotated data is scarce. Future research should optimize strategies to balance specialization with generalization through advanced pretraining methods [49], adaptive fine-tuning techniques [50], and multi-domain training frameworks [51] to maximize pretrained model utility across diverse and challenging genomic domains.

Despite these advancements, there remains significant room for improving genomic language models. One promising avenue is the development of smarter tokenizers. A guided tokenizer that incorporates biological insight into its tokenization process could substantially enhance model performance. For example, a tokenizer capable of segmenting DNA sequences into biologically meaningful units or contextually relevant patterns may improve the model’s ability to capture complex genomic information. Further research could also explore hybrid approaches that combine the strengths of different architectures or tokenization methods. Advances in pretraining strategies, such as multi-domain pretraining or biologically informed masking techniques [52], could further optimize genomic language models for diverse and challenging tasks.

## 4 Data availability

The `seqLens` models and the datasets we used in this study are available at https://huggingface.co/omicseye. The scripts that we used to get the data from the NCBI’s website, data preparation, model training, benchmarking, and visualizations are also available at https://github.com/omicsEye/seqLens.

## 5 Acknowledgments

This work was supported by the National Science Foundation under grant no. 2109688 to A.R. and K.A.C. We thank Ali R. Taheriyoun for the feedback on the manuscript text, and Ajinkya Patil and Matthew Mollerus for their assistance with the applications data.

## 6 Author contributions

M.B. and A.R. conceived the study and designed the models. M.B., A.R., and K.A.C. developed the analytical research framework. A.R. and K.A.C. supervised the study’s execution and guided data interpretation and result structuring. M.B. implemented and evaluated the models. All authors contributed to discussing the results. M.B. and A.R. wrote the manuscript. B.M. and K.A.C. reviewed and revised it, providing critical feedback that shaped the final version.

## 8 Methods

Developing advanced genomic language models requires a multifaceted approach encompassing several critical methodological components. In this manuscript, we present a comprehensive framework for creating and optimizing masked language models specifically tailored for DNA sequence analysis. Our methodology addresses key challenges in computational genomics through a systematic exploration of several interconnected research dimensions:

### 1. Data Preparation and Curation

We meticulously gathered and preprocessed diverse DNA sequence datasets to ensure robust and representative training and evaluation.

### 2. Tokenization Strategies

Recognizing the fundamental importance of sequence representation, we conducted an in-depth investigation of BPE tokenization, exploring various vocabulary sizes and their implications for model performance.

### 3. Model Architecture Design

We explored transformer-based architectures and assessed their capabilities for genomic language models.

### 4. Vector Representation Techniques

We explored different methods for representing DNA sequences as dense vector spaces, including:

- Examining different pooling strategies (CLS token, mean pooling, max pooling)
- Analyzing vector transformations using Uniform Manifold Approximation and Projection (UMAP)
- Quantitatively assessing representation quality through clustering metrics

### 5. Fine-Tuning Methodologies

We implemented sophisticated adaptation techniques to enhance model performance on specific downstream tasks:

- Low-Rank Adaptation (LoRA) for parameter-efficient fine-tuning [46]
- Domain adaptation through continual pretraining
- Innovative approaches to aggregating last hidden state vectors for classification tasks

In the subsequent sections, we provide a detailed exposition of each of these methodological components, offering insights into our approach to developing state-of-the-art genomic language models.

### 8.1 Training dataset

For pretraining and validation, we curated a dataset comprising all 19,551 representative genomes sourced from the National Center for Biotechnology Information (NCBI) database by February 23, 2024, including 18,268 bacterial, 647 archaeal, 577 fungal, 40 viral, and 1 human reference genome. Then, to create samples, we created chunks of 3200 long nucleotide sequences with 100 bases overlapping. The resulting train set has 27,831,882 samples, the validation set has 3,478,985, and the test set has 3,478,986 samples.

For the second dataset, which is a more diverse dataset in terms of the tree of life, we extended the multi-species pretraining dataset that was collected in the Nucleotide Transformers [24] study and extended it with 499 archaea genomes and 5 eukaryotic genomes (see section 4).

### 8.2 Tokenization

Tokenization is the first step in language modeling [29]. In language models, a given text sequence is split into tokens (tokenization), and then the model processes these tokens as data units. In natural language, these tokens can be sequences of characters, including complete words, punctuation marks, or even subwords. However, there are no specific words or separators in analyzing DNA sequences, so not all the natural language approaches are applicable. We used BPE [31] as a tokenization method to optimize sequence encoding. This method is more robust to small changes in DNA sequences, as reads can come from different parts of a sequence or mutations can happen only on a single base pair [3]. In the DNABERT-2 study [3], the authors provide examples of the superiority of this method to the traditional *k* -mer approach.

We trained 5 different tokenizers with vocabulary sizes of 4096, 8192, 16384, 32768, and 65536. Our choice of vocabulary size was informed by DNABERT2 [3] where they incrementally increased vocabulary size and found optimal performance at 4,096 tokens. We sampled 1 million sequences from the training dataset to train the tokenizers. Since the tokenizers have different sets of tokens and more vocabulary leads to longer tokens, tokenization results in a sequence of tokens of different lengths.

### 8.3 Model architecture

#### 8.3.1 Masked language modeling

Masked language modeling (MLM) is a technique used in natural language processing to train language models by predicting the most likely tokens at masked positions within a sequence [26, 44]. In MLM, a portion of the input tokens is replaced with a special [MASK] token, and the model is tasked with predicting the original tokens based on the context provided by the remaining, unmasked tokens. For example, given an input training sequence of *n* tokens *S* = {*t*_1_, …, *t*_*n*_}, 15% of these tokens are randomly selected and masked. The training objective of the model M is to predict these masked tokens. Each prediction yields a probability distribution over the model’s vocabularies of the model and the training objective is to minimize the cross-entropy loss between the predicted probability distribution and the true distribution of the masked tokens. This approach allows the model to learn bidirectional sequence representations, capturing dependencies between tokens in both forward and backward directions [26, 44]. MLM has proven particularly effective in domains where sequential data is prevalent, such as protein research, where sequences of amino acids are treated as words, and genomics, where sequences of nucleotides can similarly be processed as sentences composed of *k* -mers [24, 27, 28]. By leveraging MLM, language models can generate rich, context-aware embeddings that significantly enhance the understanding and analysis of complex biological sequences [3, 24].

Given a training sequence *S* = {*t*_1_, …, *t*_*n*_}, let ℳ ⊂ {1, …, *n*} denote the set of indices of the masked tokens, representing 15% of the sequence. The model *M* predicts the probability distribution over the vocabulary for each masked token *t*_*i*_, where *i* ∈ ℳ:

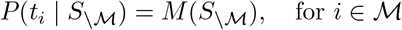

Here, *S*_*\ℳ*_ is the input sequence with the masked tokens replaced by special mask tokens. The training objective is to minimize the cross-entropy loss between the predicted probability distribution *P* (*t*_*i*_ |*S*_*\M*_) and the true distribution *t*_*i*_ for each masked token:

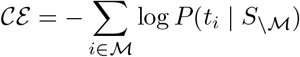

This loss function sums over the *log* probabilities of the true tokens at each masked position, which the model attempts to maximize.

#### 8.3.2 Language model architectures

For the architectures of the base model, we selected Decoding-enhanced BERT with disentangled attention (DeBERTa-v2) model architecture [26] and Evolutionary Scale Modeling (ESM) with rotary positional encoding [53]. DeBERTa-v2 is a state-of-the-art model developed by Microsoft that uses a disentangled attention mechanism and enhanced masked decoder. DeBERTa introduces a disentangled attention mechanism that separates the attention to content and position information. Unlike BERT, which represents each word with a single vector combining content and position embeddings, DeBERTa uses two separate vectors to encode a word’s content and position. The attention weights are calculated using disentangled matrices based on content and relative positions of words, reflecting that word (token) dependencies rely on both content and proximity. This mechanism captures the stronger dependency between closely related words (tokens) compared to words in different contexts and better understands the input’s positional relationships and content dependencies. The enhanced mask decoder in DeBERTa-v2 also tries to improve the training efficiency and model performance. It allows the model to predict masked tokens during the pretraining phase better. Also, DeBERTa-v2 employs parameter-sharing techniques across different layers of the network. This reduces the overall number of parameters, making the model more efficient while maintaining high performance.

In DeBERTa, the calculation of the cross-attention score between tokens *i* and *j* involves both their content and relative positional information. Specifically, *H*_*i*_ and *P*_*i*|*j*_ represent the content and the relative positional embedding of token *i* with respect to token *j*, while *H*_*j*_ and *P*_*j*|*i*_ represent the corresponding features for token *j*. The cross-attention score between tokens *i* and *j* is decomposed into four components as follows:

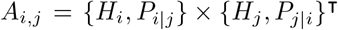

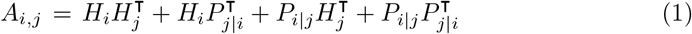

This decomposition consists of four components:

1. **Content-to-Content:** 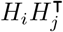, which captures the interaction between the content features of tokens *i* and *j*.
2. **Content-to-Position:** 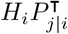, which models how the content of token *i* interacts with the relative position of token *j*.
3. **Position-to-Content:** 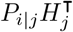, which captures the interaction between the relative position of token *i* and the content of token *j*.
4. **Position-to-Position:** 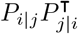, which encodes the interaction between the relative positional features of tokens *i* and *j*.

These four components work together to provide a richer representation of the relationships between tokens by incorporating both their content and relative positions [26].

Relative position encoding can be implemented by using only the *Content-to-Content* and *Content-to-Position* terms from Equation 1. However, as proposed in DeBERTa [26], including the *Position-to-Content* term is helpful to the model for fully capturing the interactions between token content and their relative positions, ensuring a more comprehensive modeling of attention weights. The same study proposed that with the first three terms of Equation 1, there is no need to include the fourth term [26].

The ESM models have two versions, and we are implementing the architecture of the ESM-2. These models are mainly developed for protein sequences, variant effect prediction, and protein structure prediction, and their main difference from other BERT family models is the ways that these models pass the last layer of the BERT encoder block to folding or classification heads of the model [27, 28]. ESM-1 used a BERT-style transformer architecture with absolute positional encodings, which were either static or learned, to provide token position information [27]. In ESM-2, the architecture remains transformer-based but introduces Rotary Position Embeddings (RoPE) [54] instead of absolute positional encodings. RoPE allows the model to extrapolate beyond the context window it was trained on. While RoPE increases the computational cost by multiplying query and key vectors with sinusoidal embeddings, it improves model quality in smaller models [28]. However, it has been reported that the performance gains diminish as model size and training time increase [28]. Here we present a refined explanation with proper notation, dimensions, and structure:

The Nucleotide Transformers v2 models adopt the architecture of the ESM-2 model, with a key modification in the feed-forward layer. In the original ESM-2 implementation, the feed-forward layer is defined as follows [55]:

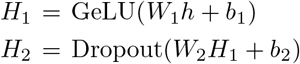

Where *W*_1_ ∈ ℝ^hidden size*×*intermediate size^, *b*_1_ ∈ ℝ^intermediate size^, *W*_2_ ∈ ℝ^intermediate size*×*hidden size^, *b*_2_ ∈ R^hidden size^, *h* ∈ ℝ^hidden size^ represents the input to the feed-forward layer, and *H*_2_ is the output. This design enables intermediate feature transformation through a higher-dimensional space defined by the *intermediate size* parameter, enhancing the model’s capacity to learn complex representations. The first transformation applies a linear projection followed by a *GeLU* activation and the second transformation applies another linear projection with a dropout layer. However, in the Nucleotide Transformers v2 feed-forward layer is as follows [56]:

1. A linear transformation:

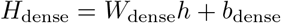

where *W*_dense_ ∈ ℝ^hidden size*×*2·intermediate size^, *b*_dense_ ∈ ℝ^2·intermediate size^.
2. Split the output to apply a gated linear unit (*GLU*) followed by a *SiLU* activation function:

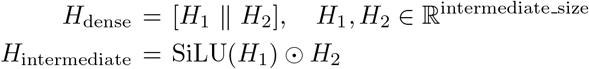

where *H*_dense_ is split along the last dimension into two equal parts, *H*_1_ and *H*_2_ and ⊙ represents element-wise multiplication.
3. The *H*_intermediate_ is then passed through a linear layer with dropout and a residual connection to *h*. In this paper, the models based on the ESM architecture share the same implementation as the Nucleotide Transformers v2 in terms of the model architecture.

Similar to BERT, DeBERTa, and ESM both use MLM for pretraining, predicting masked words based on their surrounding context. For codes and available trained models, please see the data availability section.

Since we are sharing the `seqLens` model weights, the following naming convention was adopted in this study: `seqLens_TOKENIZER-SIZE_CONTEXT-LENGTH_MODEL-SIZE`.

- Models that implement disentangled attention include a `-at` suffix.
- Models derived directly from the DeBERTa-v2 architecture (`xsmall`, `small`, and `base`) retain their size as a suffix in the model ID.
- Models based on the `ESM` approach include `esm` in their IDs.

This consistent naming convention ensures clarity, traceability, and reproducibility when sharing and referencing the model weights.

### 8.4 Model training

We initialized the model weights for each DeBERTa configuration, exploring various combinations of hidden sizes, pooler sizes, attention heads, number of layers, intermediate sizes (feed-forward layers), learning rates, and batch sizes. To evaluate the impact of relative positional encoding, we trained the model under two configurations for some setups: one using the first three terms of Equation 1 with relative positional encoding and the other without relative positional encoding and relative position attention. This resulted in the investigation of 48 models with parameters ranging from 15M to 111M. Full details of the model configurations are provided in Appendix B. We also trained the `seqLens_4096_512_89M-at-base-multi` model, using the second pretraining dataset with context length 512, batch size 1024, and learning rate of 1e-4. We trained this model for 300K which lead to roughly 150B training tokens. For the ESM models, we adhered to the architectures of the Nucleotide transformers v2 50M and 100M models, using the 4096 tokenizer and max input length of 512, batch size of 1024, and learning rate of 5e-4.

For each model (combinations of architecture, tokenizers, and parameters), we pretrained using MLM approach, where we sampled a few tokens in each sequence for masking with a probability of 15%. Of the masked tokens, 80% were replaced with the tokenizer’s mask token ([MASK]), 10% were substituted with a random word, and the remaining 10% were left unchanged. The input size for each model was 512 tokens ([CLS] + 510 + [EOS]). Each model was scheduled for 150,000 steps of training, though some failed to converge due to hardware issues, inadequate learning rates, or other unknown reasons Appendix B. We used the AdamW optimizer with parameters *β*_1_ = 0.9, *β*_2_ = 0.999, *ϵ* = 1 *×* 10^*−*8^, and a weight decay of 0.1. We applied *L*_2_ regularization only to the attention layers to improve the training process. Each training step consisted of 512, 1024, or 2048 samples (see Appendix B), resulting in approximately 768 million samples (about 3 epochs, batch size of 512) in total. We used a cosine learning rate scheduler with 5% warm-up steps (7,500 steps) and learning rates of 5e-4, 1e-4, and 5e-5. For training, we utilized a cluster of 8 Nvidia Tesla V100 SXM2 16GB GPUs (see data and code availability section). To load the training data, we used a buffer size of five times the accumulation step to avoid repeated data and introduce more randomization into the training process. We evaluated the model after each 1000 steps with 800 batches of the validation set.

### 8.5 Effect of BPE tokenizer

To evaluate the impact of tokenization methods on model performance, we conducted comparative experiments using the Nucleotide Transformer multi-species models with 50M and 100M parameters, denoted as `NT-50` and `NT-100`, respectively [24, 56]. We reinitialized the Nucleotide Transformers multi-species v2 models (`v2-50M` and `v2-100M`) and trained them from scratch using two different tokenizers: our custom-trained BPE tokenizer with a vocabulary size of 4096 and the original fixed-length *6* -mer tokenizer from the Nucleotide Transformers study [24]. Both models were trained under identical conditions, including the same parameter configuration, training and validation datasets, hyperparameters as previously outlined (see subsection 8.1), and computational resources (GPU nodes). For training, we used a batch size of 512 and monitored training loss, evaluation loss, and model performance in terms of computational efficiency. We evaluated the models by measuring the loss on the validation set, the total floating point operations (TFLOP), and the time per iteration.

### 8.6 Benchmarking datasets

#### 8.6.1 Genome benchmark dataset

The *nucleotide transformer downstream tasks* dataset represents a comprehensive genomics benchmark comprising 18 downstream tasks, exclusively derived from human samples [57]. The dataset features a mix of binary and multi-class classification tasks, carefully curated from high-quality sources. It offers diverse genomic element classifications across various scales, ranging from 300bp to 1kb sequence lengths, covering histone marks, enhancer elements, promoter regions, and splice sites. We are using the revised version of the dataset that addresses previous methodological limitations by replacing randomly generated negative samples with existing genomic sequences and implementing proper chromosome held-out test sets. The tasks include classifications of histone modifications, enhancer types (tissue-specific and tissue-invariant), promoter elements (with and without TATA-box motifs), and splice site annotations, providing a robust and consistent framework for genomic machine learning research.

#### 8.6.2 Genome origin datasets

To extend the benchmarking beyond the human genome, we prepared two additional datasets: one for plasmid/chromosome detection and another for the classification of short reads into four categories: prokaryote, eukaryote, archaea and viruses.

The `plasmid detection` dataset consists of a balanced sample of 100,000 sequences, split into 95,000 for training and 5,000 for testing. To generate these reads, we utilized the plasmid and chromosome database introduced in the DeepPlasmid study [58], which can be accessed here.

For simulating reads from the contigs, we employed the `SimLoRD` read simulation tool [59] with default parameters and a target read length of 300 base pairs (bp). The command used was:

~~~
simlord --read-reference “$input_file” \
-n “$number_of_samples” \
-fl “$read_length” \ out/”$output_file” \
--no-sam --without-ns
~~~

The `four kingdoms` dataset focuses on classifying short reads as prokaryote, eukaryote, archaea, and virus. For this dataset, we simulated reads using reference genomes of representative species from each domain, as well as the human reference genome, to ensure coverage of diverse genomic structures. The reads were simulated with a length of 150 bp using the `ART` read simulator [60]. The simulation command was:

~~~
art -ss HS25 -i “$input_genome” -o “$output_simulated” \
-l 150 -f “$fold_coverage” -na --rndSeed 1 \
--quiet -p -m 400 -s 60
~~~

This dataset is also balanced, containing a total of 80,000 samples. We divided the data into 75,000 samples for training and 5,000 for testing.

### 8.7 Benchmark

To perform extensive benchmarking, we conducted a 10-fold cross-validation using the training set of each dataset. To ensure reproducibility and consistent fold generation, we set a random seed to control for equal comparison. We reported the best performance for each fold based on the validation Matthews Correlation Coefficient (`MCC`). After completing the cross-validation, the model with the best validation performance was selected to evaluate on the test set. In addition to accuracy metrics, we also measured the evaluation speed in terms of “samples per second” and “steps per second” to compare the computational efficiency of the models.

For the benchmarking, we utilized two Nucleotide Transformers multi-species models, `v2-50M` and `v2-100M`. This selection was driven by our computational resources and the fact that, at the time of initiating this project, these models demonstrated strong performance in benchmarking tasks performed by the developers, often surpassing other implementations. We also included two ESM models, which we trained using our training dataset and a custom tokenizer with a vocabulary size of 4096. For models based on the DeBERTa architecture, where we explored the impact of various hyperparameters (such as learning rate, batch size, and the use of relative positional encoding), we selected the configuration with the lowest validation loss for inclusion in the benchmarking study. For instance, if we trained a model *M*_1_ with two different learning rates, *lr* ∈ {*lr*_1_, *lr*_2_}, only the version with the lower validation loss was used in the final evaluation. Additionally, we included models with continual pretraining (see subsection 8.11) and models with various classification heads (see subsection 8.10) to assess the impact of these modifications on performance.

### 8.8 16S gene analysis

We gathered 200 16S rRNA sequences from 9 genera within the family *Enterobacteriaceae*, including *Buttiauxella* (n=10), *Citrobacter* (n=43), *Cronobacter* (n=18), *Enterobacter* (n=58), *Escherichia* (n=10), *Klebsiella* (n=32), *Kosakonia* (n=14), *Lelliottia* (n=13), and *Raoultella* (n=2).

Sequences were aligned using MUSCLE (v3.8.31) with default high-accuracy settings [61]. Ambiguous regions containing gaps or poorly aligned segments were removed using Gblocks (v0.91b) with a minimum block length 10, no gap positions allowed, non-conserved contiguous segments longer than 8 removed, and requiring 85% sequences for a flank position [62]. Maximum likelihood phylogenetic trees were reconstructed with PhyML (v3.1/3.0 aLRT) using the HKY85 substitution model with an estimated proportion of invariant sites (0.824) and four gamma-distributed rate categories; the gamma shape parameter was estimated from the data (0.364). Internal branch reliability was assessed using SH-like aLRT support [63]. We used the online tool https://www.phylogeny.fr/ for the analysis [64].

For each sequence, we used max pooling to extract the embeddings using the `89M-at-base-multi`, `NT-100`, and `NT-50` models. Hierarchical clustering of embeddings was performed using UPGMA (average linkage) to generate an embedding dendrogram. t-SNE projections were also computed to visualize embedding clustering. Clustering agreement between genera and k-means clusters on embeddings was quantified using Normalized Mutual Information (NMI) and Adjusted Rand Score (ARS).

Pairwise cosine distances were computed between embeddings, and patristic distances were extracted from the phylogenetic tree. Mantel correlations (Spearman) were computed between embedding and patristic distance matrices using 999 permutations [34]. A null model using randomly permuted embeddings was also tested.

Tokenization of 16S sequences was performed to identify the top 15 most frequent tokens. Token frequencies were compared within the 200-sequence dataset and against the full MIMt 16S database [65]. Relative frequencies per genus were calculated by normalizing counts by the total number of tokens per genus.

For each genus, the lowest common ancestor (LCA) of all member tips was determined in both the reference phylogeny and the embedding dendrogram. A genus was considered monophyletic if the set of tips under the LCA exactly matched the set of genus members, and non-monophyletic if additional tips from other genera were included.

### 8.9 Low-rank adaptation

Fine-tuning large pretrained models for downstream tasks can be computationally intensive and financially prohibitive, often comparable to the original pretraining process [46, 66]. To address this challenge, we implemented low-rank adaptation (LoRA), an efficient fine-tuning technique for large pre-trained models [46]. Traditional fine-tuning involves updating all the parameters of a model, which can be computationally expensive and memory-intensive, especially for large-scale models. LoRA addresses this challenge by freezing the pretrained model’s original weights and introducing small, trainable low-rank matrices to specific layers. During training, only these low-rank matrices are updated which results in a significant reduction of the number of trainable parameters while maintaining the model’s performance on downstream tasks. LoRA is particularly well-suited for resource-constrained environments and applications requiring frequent fine-tuning across diverse tasks. It also reduces storage overhead since only the low-rank matrices need to be saved, making it easier to switch between tasks without retraining the entire model. Furthermore, by adding a negligible computational overhead, LoRA enables fine-tuning large models with limited hardware.

Here we investigated the effect of fine-tuning with LoRA by employing a rank of 8 and a scaling factor of 32. We targeted the query and value modules of the attention mechanism, as these are critical components for model performance and efficiency. For our experiments, we utilized two models: `NT-100` and `seqLens_4096_512_89M-at-base`. After integrating LoRA into these models, we adopted the same benchmarking setup described in subsection 8.7, ensuring consistency in evaluation metrics, datasets, and hyperparameter configurations. This allowed us to systematically assess the impact of LoRA fine-tuning on task performance, scalability, and computational efficiency.

### 8.10 Effect of different classification heads

A common approach for fine-tuning models in the BERT family for classification or regression tasks involves utilizing the `[CLS]` hidden state from the final layer. This `[CLS]` token is then passed through a feed-forward layer to make predictions [67]. A comprehensive study investigating various fine-tuning strategies for BERT models examined the utility of outputs from different layers and concluded that the last layer is generally the most informative for such tasks [45]. We extended our investigation by experimenting with the hidden states of the final layer for all non-padding tokens. Specifically, we explored three alternative strategies to using the `[CLS]` token: 1) mean pooling, 2) max pooling, and 3) concatenating the `[CLS]` token with the mean and max values (`CLS|Mean|Max`). Approaches 1 and 2 are widely used in sentence embedding applications and have been shown to outperform using only the `[CLS]` token in many scenarios [38, 39], motivating us to investigate their applicability in genomic language models. These approaches aim to capture richer contextual representations from the model.

For this analysis, we selected the `NT-50` model and implemented three different classification heads. In classification tasks, the typical approach involves using the `[CLS]` token from the final layer of the transformer as a summary representation of the input sequence. This token is then passed to a classification head, which performs a mapping from the hidden size *d* to the number of classes.

#### Mean pooling

In this approach, we compute the average of all non-padding token embeddings in the final layer. Let **H** = **h**_1_, **h**_2_, …, **h**_*n*_ be the set of hidden state vectors for each token, where **h**_*i*_ ∈ ℝ^*d*^ and *d* is the hidden size, and *n* is the number of input tokens. The average pooling is defined as:

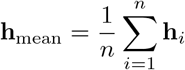

This mean vector **h**_mean_ is then fed into the classification head.

#### Max pooling

Here, we take the maximum value across all non-padding token embeddings in the final layer for each hidden dimension. The max pooling operation is defined as:

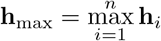

where the maximum is computed element-wise across the hidden size dimensions. The resulting vector **h**_max_ is used as the input to the classification head.

#### Concatenated Head ([CLS] — Mean — Max)

In this approach, we concatenate three different representations—the `[CLS]` token, the average-pooled vector (excluding the `[CLS]` token), and the max-pooled vector—into a single vector (`Mean` and `Max` are based on non-padding tokens). If **h**_CLS_ is the hidden state of the `[CLS]` token, the combined input to the classification head is:

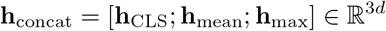

where [·; ·; ·] denotes vector concatenation. In this case, the input size to the classification head is 3 *× d*, accounting for the combined dimensions.

Each of these classification heads was implemented to assess its impact on the performance of the model during benchmarking. The results were then compared to understand how the choice of classification head influences the model’s classification performance metrics (see subsection 2.8).

### 8.11 Continual domain adaptive pretraining

To measure the effect of continual pretraining, we selected one of the models, `seqLens_4096_512_46M`, and continued pretraining as follows. We began by loading the pretrained weights while re-initializing the cosine learning rate scheduler, treating it as if the training was starting from scratch.

To evaluate the impact of this continual pretraining on model performance in benchmarking tasks, we generated two datasets using `SimLoRD` [59]. The first dataset consisted exclusively of eukaryotic samples, comprising 600K samples from the human genome, 600K samples from mouse genome (*Mus musculus*), and samples from fungal genomes (a total of 7,280,000 training samples). The second dataset contained only prokaryotic and archaeal samples, including 20M samples from 2000 bacterial and 1M samples from 100 archaeal species (a total of 20,950,000 samples).

We continued pretraining the selected model on these two datasets for 150K steps and also conducted another continuation of pretraining where we used the original training dataset that the model was trained on. In this second scenario, the learning rate scheduler was re-initialized at the start of the process, but otherwise, the training method remained unchanged. Since the learning rate is initialized, we have two separate 150K steps of training, and this is different from training with 300K steps. After completing training, we included the three resulting models in the benchmarking analysis (see subsection 8.7).

## Appendix A Tokenizers

Here, we present the information on the trained tokenizers and the tokenization comparison study.

**Fig. A1:**
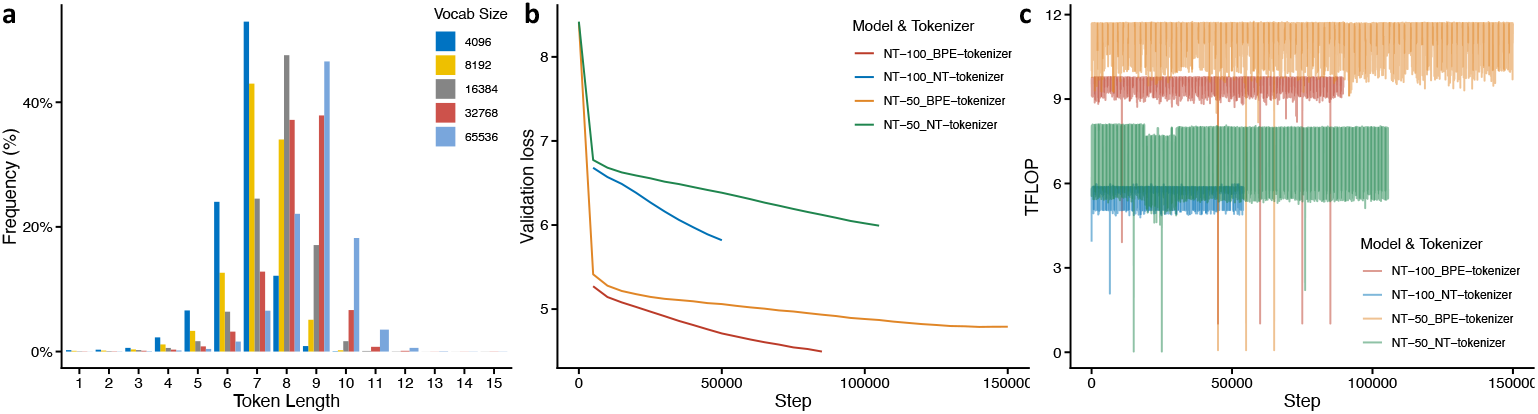
Performance and Efficiency Analysis of Tokenizers. **a**, token frequency distribution across varying vocabulary sizes, showing the impact of different token lengths on representation; **b**, validation loss comparison between models using the BPE tokenizer and the fixed-length *6* -mer tokenizer, demonstrating faster convergence and improved performance with the BPE tokenizer; **c**, computational efficiency measured in TFLOP for the BPE and *6* -mer tokenizers, highlighting the higher computational demand of the BPE tokenizer due to its more sophisticated implementation.

**Table A1:** Summary of tokenization statistics across different vocabulary sizes, showing the maximum token length, the most frequent token length with their frequencies in the vocabulary set of the tokenizer, and average token length of tokenized inputs. For example, token length 7 is the most frequent length in the tokenizer with 4,096 vocabularies and this token length has 52.93% frequency in the vocabulary set. On average, the tokenized input lengths for the 4,096 tokenizer is 4.9 characters per token.

| Vocabulary Size | Max Token Length (Frequency) | Most Frequent Token Length (Frequency) | Average Token Length |
| --- | --- | --- | --- |
| 4,096 | 10 (0.05%) | 7 (52.93%) | 4.9 |
| 8,192 | 20 (0.01%) | 7 (42.98%) | 5.3 |
| 16,384 | 21 (0.01%) | 8 (47.55%) | 5.6 |
| 32,768 | 25 (0.003%) | 9 (37.88%) | 6.0 |
| 65,536 | 40 (0.003%) | 9 (46.54%) | 6.4 |

## Appendix B Model pretraining

We initialized a set of models with varying parameter sizes for each tokenizer. For example, each tokenizer was used to create a model configuration with 4 layers, 4 attention heads, a hidden size of 512, and an intermediate size of 2048. The number of layers, attention heads, hidden size, and intermediate size influence the architecture of a typical encoder model Figure B2a. Each model was trained for 150K steps using batch sizes of 256, 512, and 1024, learning rates of 5e-4, 1e-4, 5e-5, and 5e-1, and different attention mechanisms (Table B2). This resulted in training 48 different combinations of models and hyperparameters with DeBERT-v2 architecture. The runtime for the pretraining of each combination with Nvidia Tesla V100 SXM2 16GB GPUs took two to seven days.

Each model was trained using one or a subset of the combinations of the hyperparameters, as detailed in Table B2. We also investigated the `xsmall`, `small`, and `base` parameter sets that were introduced in the DeBERTa-v2 study [26]. Figure B2b shows the validation loss of these three setups with the 4096 tokenizer. The final validation loss at the end of training is closely tied to the model size, with larger models generally achieving lower losses. We validated loss for the `small` model with the 4096 tokenizer across different hyperparameter combinations (Figure B2c). Our results indicate that the best loss for the `small` model was achieved with a learning rate of 5e-4 and a batch size of 1024 samples, underscoring the importance of proper hyperparameter tuning. Figure B2d compares the training performance of two models: one with 15M parameters (left panel) and another with 46M parameters (right panel). For these models, we explored the effects of various learning rates, batch sizes, and the use of disentangled attention. Although the goal was not to comprehensively analyze the impact of disentangled attention, for these two specific models, the best setup was achieved using standard attention. However, for the `base` model with the 4096 tokenizer, the best performance was observed with the model employing disentangled attention (validation loss: 4.932), outperforming the equivalent setup without disentangled attention (validation loss: 5.287). The optimal setup for this model was a learning rate of 1e-4 and a batch size of 512. To build the `seqLens` models, we also investigated other configurations from the DeBERTa-v2 models. For a detailed list of models subsection 8.3 and Table B2). After training, the best-performing configuration for each of the model sizes (e.g., the 15M parameter model) was selected based on evaluation loss. These selected models then progressed to the next phase, which involved benchmarking their performance on downstream tasks (see subsection 8.4).

**Table B2:**
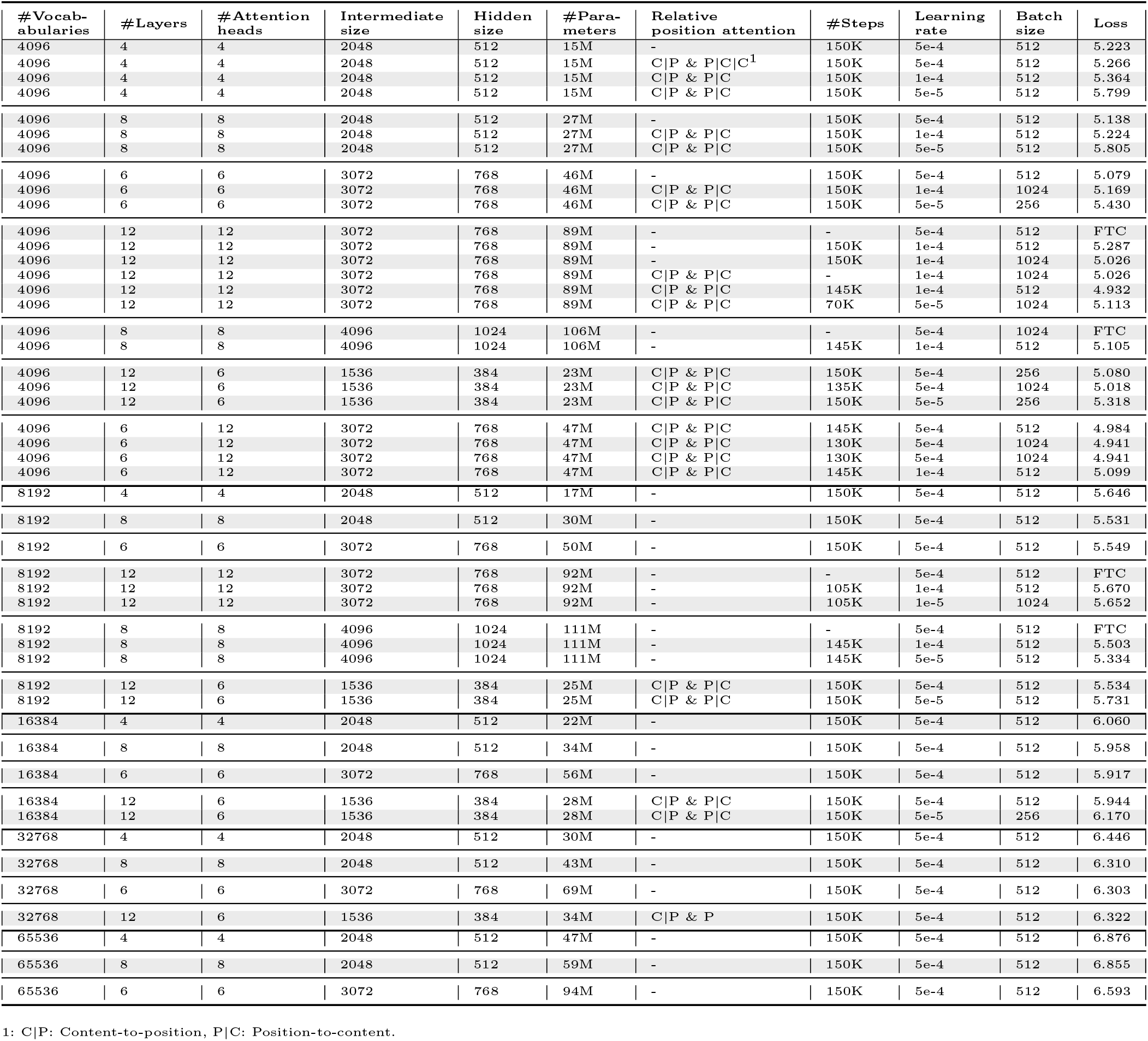
Models parameters and pretraining loss results.

**Fig. B2:**
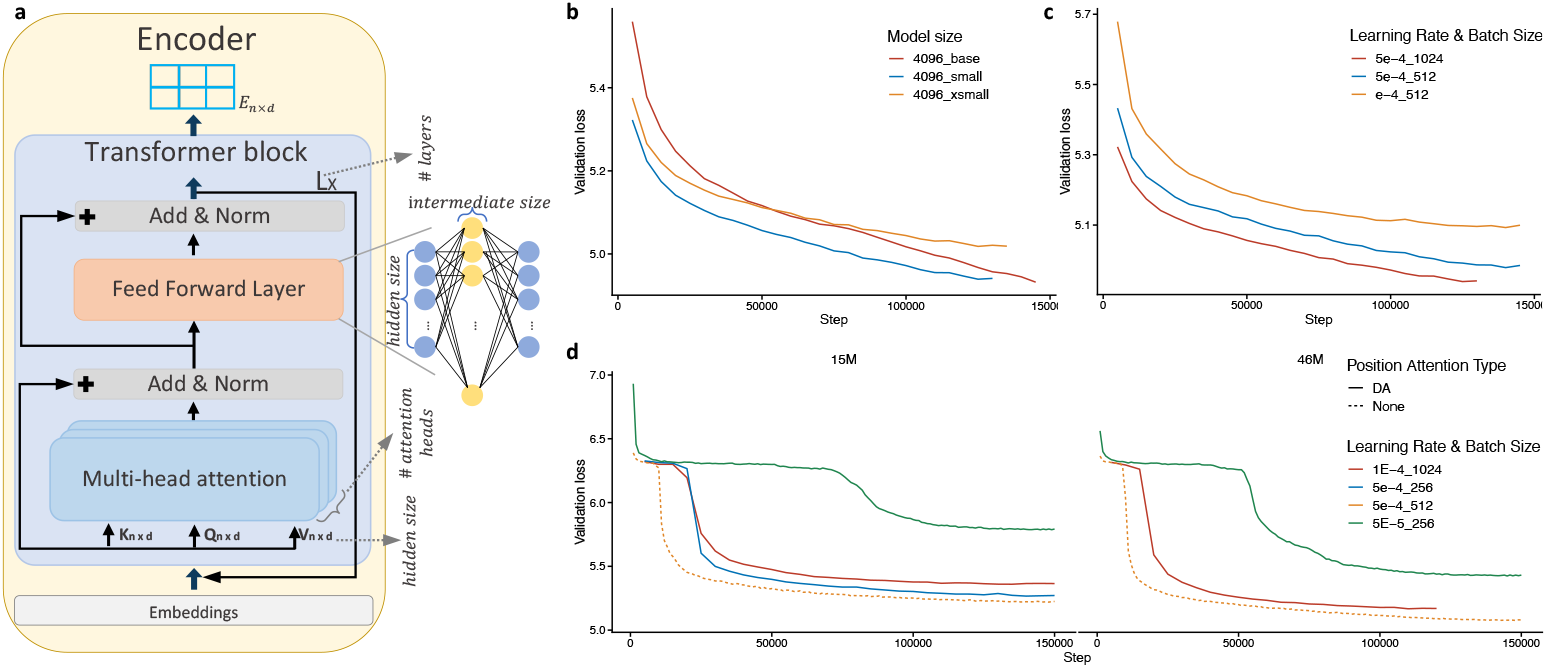
Training dynamics and hyperparameter impacts on model performance. **a**, An overview of model architecture components, illustrating how the number of layers, attention heads, hidden size, and intermediate size influence the design; **b**, Validation loss curves for the `xsmall`, `small`, and `base` configurations of the DeBERTa-v2 model using the 4096 tokenizer, showing that larger models generally achieve lower losses; **c**, Validation loss curves for the `small` model with the 4096 tokenizer, highlighting that the best performance is achieved with a learning rate of 5e-4 and a batch size of 1024; **d**, Training loss comparisons for models with 15M parameters (left) and 46M parameters (right), including variations with disentangled attention (DA).

## Appendix C Benchmarking results of models

Here, we present the results of model benchmarking on downstream tasks, including validation and test score comparisons across benchmarking datasets. These plots allow for a direct comparison of different architectures and an investigation into the impact of tokenizers with varying vocabulary sizes within the same architecture.

**Fig. C3:**
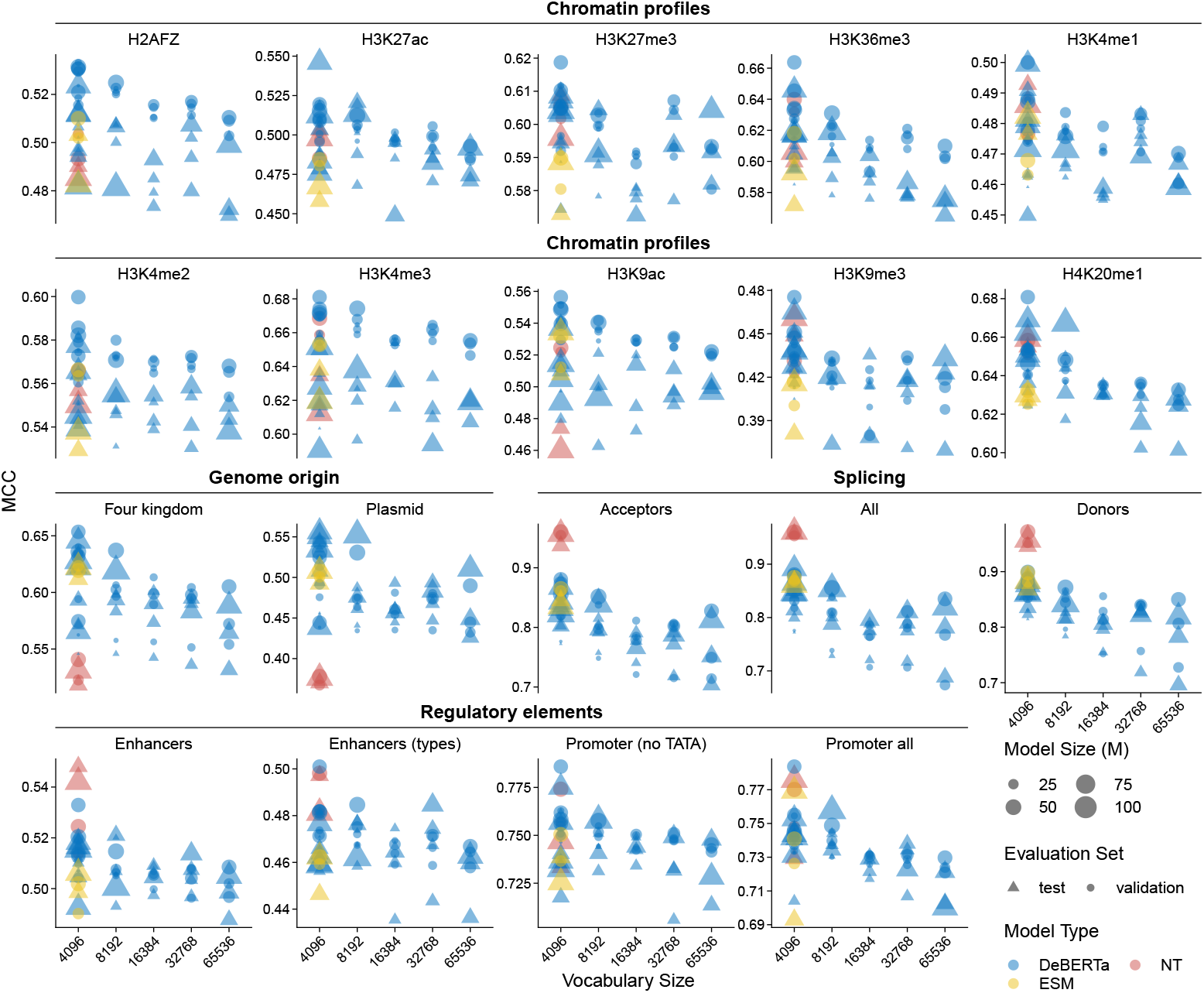
Validation and test scores of models across benchmarking datasets. The figure compares the performance of models trained in this study (based on DeBERTa and ESM architectures) with Nucleotide Transformer models (`NT-50` and `NT-100`). Validation scores are the average of the scores for each fold. Results demonstrate no universally best-performing model, as smaller models occasionally outperform larger ones across datasets. A negative correlation is observed between tokenizer size and model performance, with larger vocabularies generally leading to lower performance. Models trained on bacterial genomes in this study perform better on the `four kingdoms` and `plasmid detection` datasets, while Nucleotide Transformer models, pretrained on datasets with greater eukaryotic representation, perform better on human genome-specific tasks such as chromatin profiles.

Figure C4 and Figure C5 illustrate a comparison between models of the same architecture using tokenizers with different vocabulary sizes. The benchmarking results indicate that, in most cases, models benefit from tokenizers with smaller vocabulary sizes, leading to improved performance.

**Fig. C4:**
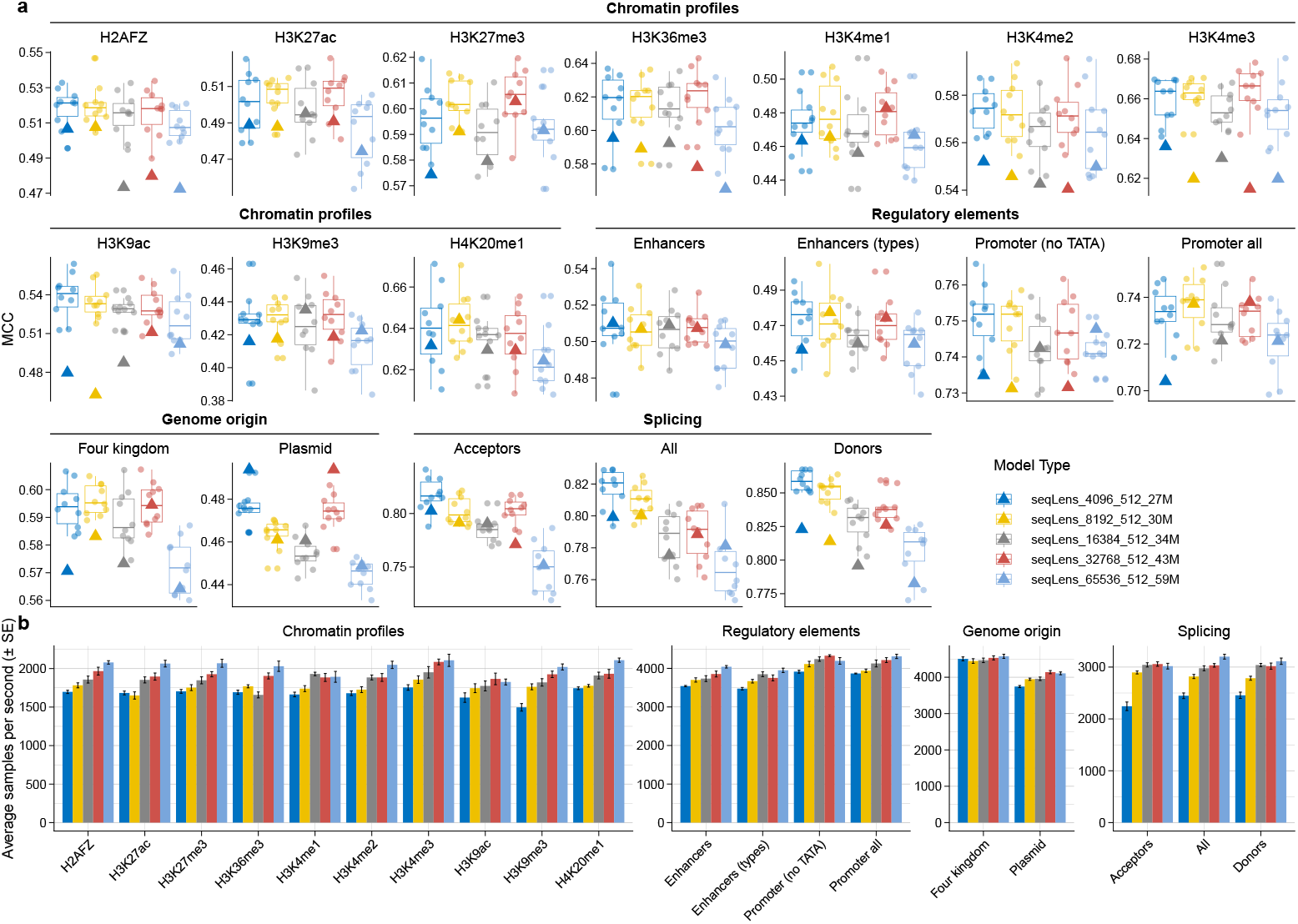
Validation and test scores of models across benchmarking datasets. **a**: The figure compares the performance of the models with the same parameter configuration but different tokenizer vocabulary sizes. Boxplots are based on the validation scores (circles) and the triangle represents the models’ performance on the test set. **b**: The figure compares the processing time of the models by comparing the average samples per second.

**Fig. C5:**
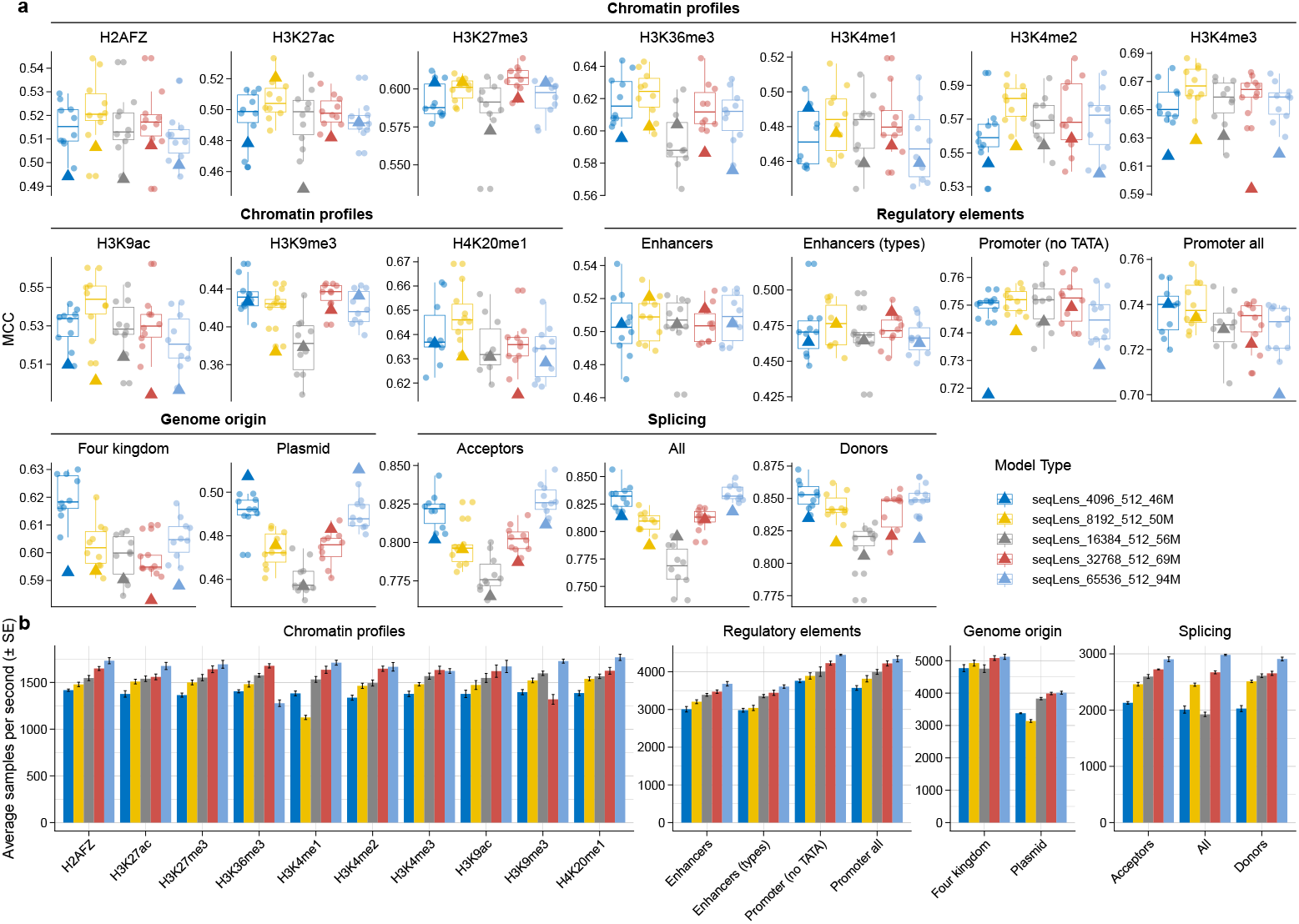
Validation and test scores of models across benchmarking datasets. **a**: The figure compares the performance of the models with the same parameter configuration but different tokenizer vocabulary sizes. Boxplots are based on the validation scores (circles) and the triangle represents the models’ performance on the test set. **b**: The figure compares the processing time of the models by comparing the average samples per second.

## Appendix D Vector representation of DNA sequences

This appendix presents a detailed analysis of the vector representations generated by the models discussed in the main text. To explore the structure and properties of these representations, we conducted a Principal Component Analysis (PCA) on the output vectors and visualized the first two principal components (PCs) through pair plots. Each figure provides a comprehensive view of how these components interact and distribute for specific datasets and summarization methods.

The analysis covers both pretrained and fine-tuned `seqLens_4096_512_89M-at-base` models across different datasets (`four kingdoms` and `plasmid detection`) and summarization methods (`[CLS]` token, mean pooling, and max pooling). These visualizations help to uncover patterns, clustering behavior, and separability in the vector spaces, highlighting the impact of fine-tuning on embedding quality and task alignment.

Encoder models output a vector of size equal to the hidden dimension *d* for each input token. If an input sequence is tokenized into *n* tokens, the model generates a vector for each of these tokens, including special tokens such as `[CLS]`. A common way to summarize information within an input sequence is to pass the sequence through the model and then use its vector representations [38, 39]. In this study, we selected the `base` model that we trained to explore its vector outputs on two datasets: `four kingdoms` and `plasmid detection`. This analysis was performed for both the pretrained model and the fine-tuned model to examine the differences in performance and vector representations after fine-tuning.

**Fig. D6:**
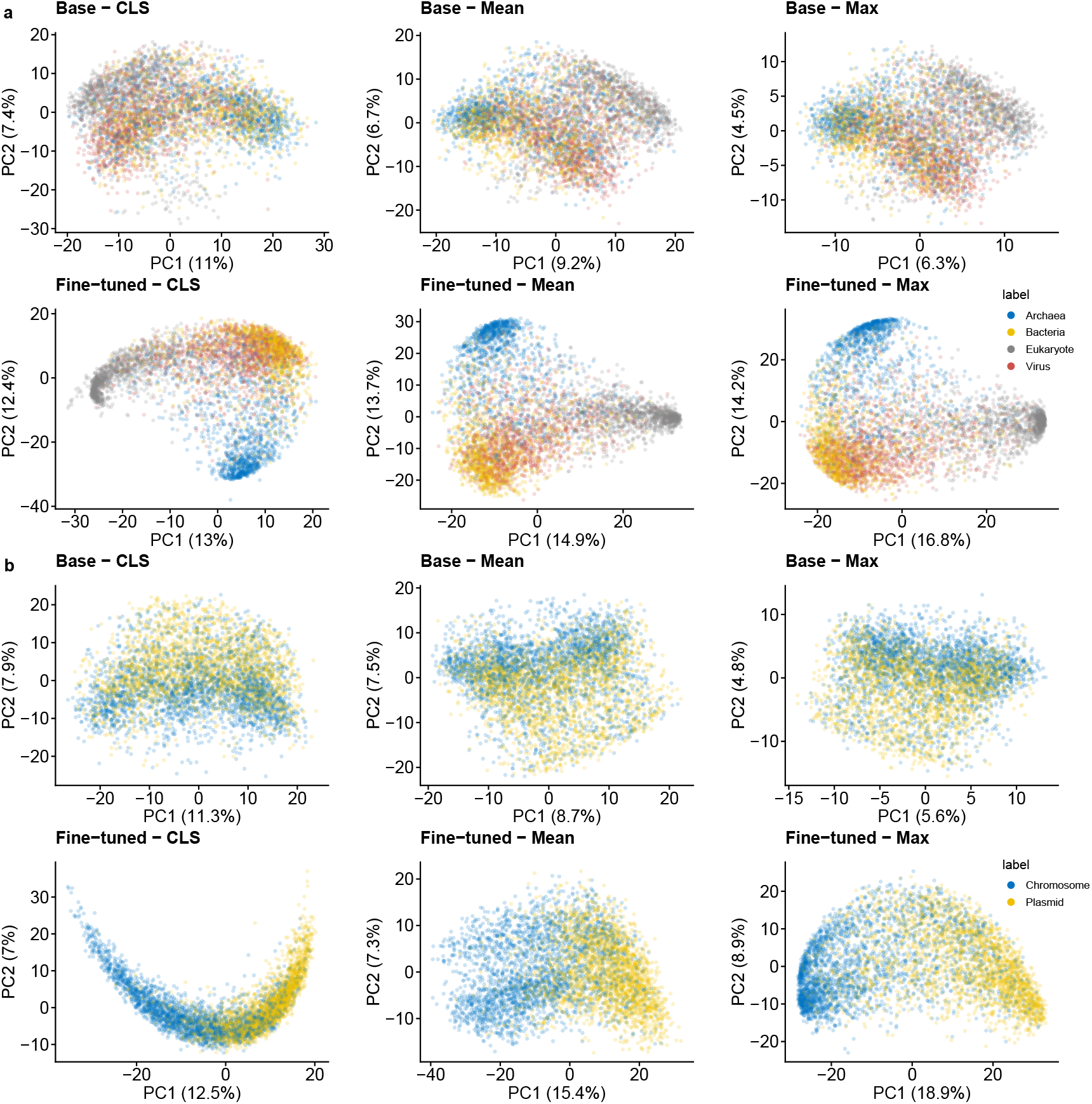
Pair plot of the first two principal components (PCs) of the vector representations from the models on the `four kingdoms` and `plasmid detection` dataset. Each panel shows the interaction between two PC 1 and PC 2, revealing patterns and clustering in the embedding space.

